# NEAT1 promotes genome stability via m6A methylation-dependent regulation of CHD4

**DOI:** 10.1101/2024.02.05.578920

**Authors:** Victoria Mamontova, Barbara Trifault, Anne-Sophie Gribling-Burrer, Patrick Bohn, Pit Preckwinkel, Werner Schmitz, Peter Gallant, Dimitrios Papadopoulos, Martin Eilers, Tony Gutschner, Redmond P. Smyth, Kaspar Burger

## Abstract

Long non-coding (lnc)RNA emerge as regulators of genome stability. The nuclear enriched abundant transcript 1 (*NEAT1*) locus encodes two lncRNA isoforms that modulate gene expression, growth and proliferation in mammals. Interestingly, NEAT1 transcripts are overexpressed in many tumours and induced by DNA damage, suggesting a genome-protective function. However, the precise role of NEAT1 in the DNA damage response (DDR) is unclear. Here, we investigate the expression, modification levels, localization and structure of NEAT1 in response to DNA double-strand breaks (DSBs) induced by the topoisomerase-II inhibitor etoposide or the locus-specific endonuclease AsiSI. We find that induction of DSBs increases both the levels and N6-methyladenosine (m6A) marks on NEAT1, which promotes alterations in NEAT1 secondary structure and accumulation of hyper-methylated NEAT1 at a subset of promoter-associated DSBs to facilitate efficient DSB signalling. The depletion of NEAT1, in turn, delays the response to DSBs and triggers elevated DNA damage. The genome-protective role of NEAT1 is mediated by the RNA methyltransferase 3 (METTL3) and involves spreading of the chromodomain helicase DNA binding protein 4 (CHD4) upon release from NEAT1. Together, we describe a novel RNA-dependent DDR pathway that couples NEAT1 to the recognition and repair of DSBs.

## INTRODUCTION

Chromosomes encode genetic information that requires faithful inheritance to maintain genome stability. The DNA damage response (DDR) is a multifaceted signalling network that recognizes and repairs DNA lesions to prevent genomic instability and tumorigenesis(1–3). Unscheduled or excessive RNA synthesis creates open chromatin and exposes DNA, which leads to highly-toxic DNA double-strand breaks (DSBs), if left unrepaired(4, 5). Thus, the DDR globally impairs transcription during DSB repair (DSBR)(6, 7). In mammals, DSBR is governed by kinases like *Ataxia-telangiectasia mutated* (ATM), which activates >100 factors to amplify the DDR and catalyse DSBR via homologous recombination (HR) or non-homologous end joining (NHEJ)(8, 9). Intriguingly, approximately 40% of DNA damage-induced phosphorylation events modify factors related to nucleic acid metabolism, in particular RNA-binding proteins (RBPs)(10, 11), suggesting a crosstalk of the DDR with RNA metabolism. Indeed, the production and processing of non-coding transcripts promotes DSBR in concert with canonical DSBR factors(12, 13). The endoribonuclease Dicer, for instance, accumulates in the nucleus and processes RNA polymerase II (RNAPII)-dependent nascent transcripts into small RNA upon induction of DSBs, which promotes the recruitment of some DSBR factors like the p53-binding protein 1 (53BP1), as shown by us(14–16) and others(17–19). DSBs can also undergo RNA-templated DNA repair or engage transcripts as scaffolds for repair factors(20–23). Thus, several modes of RNA-dependent DSBR operate in context of chromatin and the cell cycle, which complements a large body of evidence that defines DNA-binding factors as crucial components of DSBR.

Long non-coding (lnc)RNA regulate multiple cellular processes(24). Interestingly, lncRNA seed the formation of nuclear membraneless organelles, such as paraspeckles, and are also linked to genome maintenance(25, 26). For instance, we recently showed that NONO, a multifunctional RBP and core component of paraspeckles, undergoes nucleolar relocalization to stimulate DSBR by shielding aberrant transcripts from broken chromatin(27, 28). Paraspeckles are phase-separated nuclear bodies that condensate around two isoforms of the lncRNA NEAT1 (NEAT1_1 and NEAT1_2) and regulate gene expression in the interchromatin space of mammalian nuclei(29, 30). The long isoform (NEAT1_2, 23 kb) is a nuclear architectural lncRNA with modular domains and undergoes a core-shell arrangement to scaffold the association of >40 RBPs, which facilitates the retention and editing of messenger (m)RNA and amplifies micro (mi)RNA biogenesis in unperturbed cells(31–34). The short isoform (NEAT1_1, 3.7 kb) is less characterized, but has recently been identified as a stimulator of glycolysis in the cytoplasm(35). Interestingly, elevated levels of NEAT1 promote tumour development in mouse models and function as a prognostic marker for patient survival(36, 37). High levels of NEAT1, for instance, correlate with enhanced growth and proliferation, as well as poor prognosis for disease-free survival in colon cancer(38). However, NEAT1 transcripts are also tumour suppressive(39). This suggests that NEAT1 transcripts have a dual role in cancer formation, which may depend on the cell type-specific expression or the differential subcellular localization of NEAT1 isoforms. Intriguingly, the levels of NEAT1 transcripts are dynamically regulated and responsive to various kinds of stress, including DNA damage(40–44). Moreover, the depletion of NEAT1 transcripts diminishes the expression and activation of a subset of critical DSBR factors, such as CHK2, RPA32 and BRCA1(45), suggesting a role for NEAT1 in the DDR. However, the molecular principles that engage NEAT1 in genome maintenance remain poorly understood.

Here we show that NEAT1 associates with a subset of DSBs to promote genome stability in human cancer cells. NEAT1 chromatin occupancy depends on the RNA methyltransferase 3 (METTL3), which places N6-methyladenosine (m6A) marks on NEAT1 *in vitro* and enhances the association of NEAT1 with DSBs *in vivo*. The genome-protective role of NEAT1 is accompanied with changes in the secondary structure of NEAT1_1 and the release of the histone deacetylase CHD4 from hyper-methylated NEAT1. The depletion of NEAT1 transcripts, in turn, elevates DNA damage, diminishes DSB signalling efficacy and hypersensitizes cells to etoposide treatment. Together, our data suggest a role for NEAT1 in the recognition and repair of DSBs and point toward a novel RNA-dependent DDR pathway.

## MATERIALS AND METHODS

### Tissue culture

Human U2OS, AsiSI-ER expressing U2OS (DIvA, kind gift from Gaelle Legube), HEK293 and MS2-tagged HEK293:24xMS2-NEAT1 cells were cultured in Dulbecco’s modified eagle’s medium (DMEM, Gibco) with 10% fetal bovine serum (FBS, Capricorn), 100 U/mL penicillin-streptomycin (Gibco), 2 mM L-glutamine (Gibco) at 37°C and 5% CO2. Cells were incubated with etoposide (Sigma, 20 µM) for 2 h or 4-hydroxytamoxifen (4-OHT, Sigma, 10 µM) for 4 h, or preincubated with ATM inhibitor KU-55933 (Hycultec, 1 µM) for 2 h and METTL3 inhibitor STM2457 (Hycultec, 10 µM) for 16 h, respectively, unless stated differently.

### Transfection and cloning

Transfection of small-interfering (si)RNA, antisense oligonucleotides (ASOs) or plasmids pcDNA-FLAG-METTL3 and pcDNA-FLAG-METTL3-APPA (kind gifts from Alessandro Fatica), pDRGFP and pCBASceI (kind gifts from Maria Jasin), pHAGE-EFS-MCP-3XmRuby3-nls and pHAGE-EFS-MCP-sfGFP-nls (kind gifts from Ling Ling Chen), and pcDNA-FRT-TO-CHD4-GFP (kind gift from Gernot Längst) was performed using Lipofectamine 2000 (Invitrogen) and Opti-MEM (Gibco) following the manufacturer’s protocol. The CHD4-GFP mutant was cloned with selective primers (fwd, 5’-GGATGCTACAGGTGGAACCCTGCACCCCTA-3’; rev, 5’-GCCCAGGCCCGACGCCAT-3’) and a Q5 site-directed mutagenesis kit (NEB) following the manufacturer’s protocol, and verified by Sanger sequencing. The NEAT1_1 *in vitro* transcription (IVT) template was generated from pCRII-TOPO-NEAT1_1 (kind gift from Archa Fox) by double digestion with BamHI/NcoI (NEB), PCR amplification using Phusion polymerase (NEB) with selective primes (fwd, 5’-CCCAGTCACGACGTTGTAAAACG-3’; rev, 5’-GTAACGGCCGCCAGTGTG-3’), re-digestion with BamHI (NEB) and purification with a PCR clean-up kit (NEB) following the manufacturer’s protocols. For manipulation with oligonucleotides, cells were transfected (6 h) on two consecutive days with 100 nM siRNA (Table S1) or 100 nM ASOs (Table S2).

### Construction of HEK293:24xMS2-NEAT1 cells by CRISPR/Cas9

To obtain single cell clones carrying 24xMS2 stem loops at the 5’ end of the *NEAT1* gene in HEK293 cells, we transfected wild type HEK293 cells with 1.3 µg bicistronic nuclease plasmid and 0.7 µg of MS2 knock-in donor plasmid (kind gifts from Ling Ling Chen) using TurboFect transfection reagent (Thermo) as described(46). Puromycin (1µg/mL, Thermo) was added 24 h later to increase knock-in efficiency. To obtain individual knock-in clones, cells were sorted after another 48 h into 96-well plates using a BD FACSMelody Cell Sorter (BD Biosciences). Targeted knock-in of the 24xMS2 stem loops was confirmed by locus- and knock-in-specific PCR. Briefly, 100 ng genomic DNA, isolated from individual clones using the GenElute Mammalian Genomic DNA Miniprep Kit (Sigma), was mixed with DreamTaq DNA Polymerase, 10x buffer and dNTPs (all from Thermo) and one of the following junction primer pairs with the respective forward primer located outside of the left homology arm of the knock-in donor construct: MS2-NEAT1_KI_forward_1 (5’-AGGAGTTCACCAGGTTTGCTT-3’) and MS2-NEAT1_KI_reverse_1 (5’-CCCCCTCGTCTCATCTAACTC-3’); MS2-NEAT1_KI_forward_2 (5’-TCAGATGACACACAGTCACCAGTT-3’) and MS2-NEAT1_KI_reverse_2 (5’-GAGCTATCTAGATGCATGCTCGAG-3’). PCR products were separated by agarose gel electrophoresis and stained with ethidium bromide (5 µg/mL) under UV-light. Only clones with correct PCR products in both PCR reactions were considered knock-ins and used for subsequent assays.

### Fluorescence-activated cell sorting (FACS)

Cells were washed in phosphate buffered saline (PBS), trypsinized, resuspended in DMEM and centrifuged (1500 rpm, 5 min, 4°C). Pellets were washed in PBS, centrifuged (1500 rpm, 5 min, 4°C), resuspended in 1 ml PBS and fixed in 4 ml ice-cold 100% ethanol (-20°C, overnight). Cells were pelleted (1500 rpm, 10 min), washed in PBS, pelleted again and resuspended in 1 ml PBS. 1x10^6^ cells were stained with 54 µM propidium iodide (Sigma) in the presence of 24 µg/ml RNase A (Sigma) (30 min, 37°C, dark), sorted and analysed by a FACSDiva 9.0.1 flow cytometer (50.000 events per condition) and software (BD Biosciences).

### Immunoblotting and immunoprecipitation

Proteins were assessed as whole cell extracts, directly lysed, boiled and sonicated in 4x sample buffer (250 mM tris-HCl pH6.8, 8% SDS, 40% glycerol, 0.8% β-mercaptoethanol, 0.02% bromophenol blue). Samples were separated by SDS-PAGE and transferred to nitrocellulose membranes (Cytiva) stained with 0.5% ponceau S/1% acetic acid, blocked, washed in PBS/0.1% triton x-100/5% milk (PBST), probed with selective antibodies (Table S3) or a streptavidin-HRP probe (Invitrogen) and visualized with an ECL kit (Cytiva) and an imaging station (LAS-4000, Fuji or Fusion FX, Vilber) following the manufacturer’s protocols. Signals were quantified by ImageJ (NIH). For immunoprecipitation, cells were trypsinized, washed in PBS and centrifuged (1200 rpm, 5 min). Pellets were lysed (10 min on ice) in 5 volumes of IP buffer (200 mM NaCl, 0.5 mM EDTA, 20 mM HEPES, 0.2% NP-40, 10% glycerol, 100U Ribolock inhibitor, Thermo, 1x protease/phosphatase inhibitor, Roche). Lysates were centrifuged (12000 rpm, 12 min) and supernatants were incubated (2 h, 4°C) with 2-5 µg primary antibodies, pre-conjugated to 25 µL protein G dynabeads (Invitrogen). Immunocomplexes were immobilized on a magnet (Invitrogen), washed three times with 800 µL IP buffer (10 min, 4°C) and eluted with sample buffer (5 min, 95°C).

### Imaging

Cells were grown on coverslips (Roth), washed in PBS, fixed (10 min) in 3% paraformaldehyde (Sigma), washed in PBS (3 times, 5min), permeabilized with PBS/0.1% triton x-100 (10 min) and blocked with PBS/10% FBS (2 h, 4°C). Primary and secondary antibodies (Table S3) were diluted in PBS/0.15% FBS and incubated in a humidified chamber (overnight, 4°C or 2 h, RT), respectively. Cells were washed between incubations with PBS/0.1% triton x-100 (3 times, 5 min), sealed in 6-diamidino-2-phenylindole (DAPI)-containing mounting medium (Vectashield), and imaged by confocal microscopy (CLSM-Leica-SP2, 1024x1024 resolution, 63x, airy=1). Channels were acquired sequentially, between frames, with equal exposure times. Colocalization was assessed by using RGB profiler (ImageJ) and by the calculation of the Pearson correlation coefficient using JACoP (ImageJ). Proximity ligation assays (PLAs) were performed with a Duolink in-situ PLA kit (Sigma) following the manufacturer’s protocol. For RNA-PLA assays, cells were washed, fixed and permeabilized as above, then blocked with RNA-PLA blocking buffer (10 mM tris-acetate, pH7.5, 10 mM magnesium acetate, 50 mM potassium acetate, 250 mM NaCl, 0.25 µg/µL BSA, 0.05% triton x-100, 100U Ribolock inhibitor, Thermo) at 4°C for 1 h. Cells were incubated (4°C, overnight) with RNA-PLA probes (Table S4), which were pre-diluted to 100 nM in RNA-PLA blocking buffer and pre-heated at 70°C for 3 min. Samples were washed three times in PBS, blocked with custom PLA blocking buffer (2 h, 4°C), washed again three times in PBS, and incubated with appropriately diluted primary antibody (4°C, overnight). The subsequent steps were performed following the manufacturer’s protocol, but using the MINUS PLA probe only. RNA fluorescence in situ hybridization (RNA-FISH) experiments were performed following the protocol for simultaneous immunofluorescence and FISH in adherent cells from Stellaris. A panel of pre-designed validated human NEAT1 5’ segment probes with Quasar570 dye has been used for hybridizations (Stellaris, SMF-2036-1).

### Neutral comet assay

Glass slides were covered with 0.01% poly-L-lysine solution (Sigma) and 1% agarose (Roth) in ddH2O and incubated in a hybridization oven (UVP) at 70°C overnight. Cells were trypsinized, washed in PBS, counted and diluted to 1x10^5^ cells/mL. The cell suspension was mixed 1:1 with 1.5% low melting temperature agarose gel (Biozym) in PBS at 37°C, pipetted on preincubated glass slides and flattened with a coverslip. The slides were incubated for 10 min at 4°C, coverslips were removed, lysis buffer (2.5 M NaCl, 0.1 M EDTA, 0.1 M tris-HCl pH10, 1% triton x-100) was added, and slides were covered with parafilm and incubated (1h, 4°C). Slides were washed twice in PBS and subjected to electrophoresis (1 V/cm, 15 min, 4°C) in neutral comet buffer (100 mM tris base pH8.5, 300 mM sodium acetate). Slides were fixed in 70% ethanol and dried (RT, overnight), stained in PBS containing 1x SYBR gold (Thermo) for 20 min protected from light, imaged by confocal microscopy and quantified using CometScore freeware v2.0 software.

### RNA analytics

Total or immunoselected RNA was isolated using TRIzol (Invitrogen) following the manufacturer’s protocol. cDNA was synthesized using SuperScriptIII reverse transcriptase (Invitrogen) with gene-specific primers (Table S5) and quantified upon reverse transcription by quantitative PCR (RT-qPCR) in a thermocycler (Applied) with PowerUp SYBR green master mix (Applied) following the manufacturers protocols. For dot blots, total RNA was resuspended in ddH2O with 0.02% methylene blue, heated (5 min, 72°C), spotted on a nylon membrane (Cytiva), crosslinked (120 mJ/cm^2^) using a UV-crosslinker (Analytik Jena), blocked in PBS/0.1% triton x-100/0.5% SDS (20 min), washed with PBS/0.1% triton x-100 (20 min), incubated (4°C, overnight) with a selective antibody, washed with PBS/0.1% triton x-100 (20 min), and visualized with an ECL kit (Cytiva).

### RNA immunoprecipitation (RIP)

For RIP, 10 µg total RNA was diluted in 800 µL IP buffer and incubated with 10 µg selective antibody at 4°C overnight. Immune complexes were pulled down for 45 min at 4°C with 25 µL protein G dynabeads (Invitrogen), captured on a magnet and washed 4 times in 800 µL IP buffer. The immunoselected RNA was purified using TRIzol along with 1 µg total RNA input. For qualitative analysis, samples were mixed with 1 volume of 2x urea dye (7 M urea, 0.05% xylene cyanol, 0.05% bromophenol blue), incubated at 75°C for 10 min and separated for 30 min at 350 V in 1x TBE buffer (90 mM tris base, 90 mM boric acid, 2 mM EDTA) on a 6% PAGE gel with 7M urea. Gels were stained in 1x TBE buffer containing 1x SYBR gold (Thermo) for 20 min protected from light. RNA was visualized on a transilluminator (Thermo). For quantitative analysis, the amount of total and immunoselected RNA were determined by RT-qPCR.

### *In vitro* transcription (IVT), pull-downs and S-adenosyl-methionine (SAM)fluoro assay

To synthesize non-methylated (or biotin-16-UTP-labelled) NEAT1_1 i*n vitro*, 1 µL (500 ng/µL) IVT template was incubated with IVT mix (7 µL ddH2O, 2 µL 10x T7 reaction buffer, 2 µL 100 mM DTT, 2 µL 10 mM ATP/CTP/GTP mix, 2 µL 10 mM UTP (or 1.3 µL 10 mM UTP mixed with 0.7 µL 10 mM biotin-16-UTP), 2 µL T7 RNA labelling polymerase mix, 2 µL 100U Ribolock inhibitor, Thermo) from the high yield T7 biotin16 RNA labelling kit (Jena) at 37°C for 4 h. Reactions were centrifuged (13000 rpm, 5 min, 4°C) and a 10% aliquot was resuspended in 2x RNA loading buffer (50% formamide, 15% formaldehyde, 40 mM MOPS, 10 mM NaAc, 1 mM EDTA pH7.0, 0.1% bromophenol blue, 10 µg/mL ethidium bromide) to monitor the integrity of the IVT product by separation on a 1.2% agarose gel containing 5.5% paraformaldehyde and 1x MOPS buffer (40 mM MOPS, 10 mM NaAc, 1 mM EDTA pH7.0) for 90 min at 100V, and staining under UV-light. For pull-downs, 1 µg non-methylated NEAT1_1 IVT product was radio-labelled with radioactive labelling mix (1 µL 10x PNK buffer, NEB, 1 µL NEAT1_1 IVT product, 1 µL T4 PNK, NEB, 1 µL γ-^32^P-ATP, Hartmann, 1 µL 100U Ribolock inhibitor, Thermo, 5 µL ddH2O) for 40 min at 37°C. Radio-labelled NEAT1_1 was centrifuged (3200 rpm, 5 min) with G-25 columns (Cytiva), diluted in 800 µL IP buffer and incubated (2 h, RT with rotation) with endogenous CHD4 that was immobilized on 25 µL antibody-conjugated protein G dynabeads (Invitrogen) upon immunoprecipitation from whole cell lysates. Immunocomplexes were captured on a magnet, washed twice with 800 µL IP buffer and incubated (5 min, RT) in the absence or presence of 10U benzonase (Millipore). Coenriched, radio-labelled NEAT1_1 was purified by TRIzol extraction, separated on a 1.2% agarose gel containing 5.5% paraformaldehyde and 1x MOPS buffer (40 mM MOPS, 10 mM NaAc, 1 mM EDTA pH7.0) for 90 min at 100V, and visualized by autoradiography with hyperfilms (Cytiva). The biotin-16-UTP-labelled NEAT1_1 IVT product (1 µg) was immobilized on 25 µL streptavidin C1 dynabeads (Invitrogen), washed twice with 800 µL IP buffer and incubated (2 h, RT with rotation) with FLAG-METTL3 or CHD4-GFP variants that were immobilized on 25 µL antibody-conjugated protein G dynabeads (Invitrogen) upon expression in HEK293 cells and immunoprecipitation (4°C, 4 h with rotation) from whole cell lysates. Pull-down complexes were captured on a magnet, washed twice with 800 µL IP buffer,eluted by boiling (95°C, 5 min) in sample buffer and analysed by immunoblotting. For the SAMfluoro methylation assay (G-Biosciences), FLAG-METTL3 variants were immobilized on 25 µL antibody-conjugated protein G dynabeads (Invitrogen) upon expression in HEK293 cells and immunoprecipitation (4°C, 4 h with rotation) from whole cell lysates, diluted in 250 µL SAM methylation assay buffer and mixed with 1 µg non-methylated NEAT1_1 IVT acceptor substrate pre-diluted in 50 µL ddH2O containing 2 µL 100U Ribolock inhibitor (Thermo). 100 µL aliquots from the mix were put to 96-well plate format and incubated with 100 µL/well SAM methyltransferase assay master mix containing SAM methyltransferase assay buffer with additive, SAMfluoro enzyme mix, SAMflourometric mix and SAM substrate according to the manufacturer’s protocol. Samples were incubated at 37°C and Resorufin emission was determined every minute for 20 minutes with a plate reader (TECAN).

### Subcellular fractionation

Trypsinized and washed cells were lysed in 5 volumes of hypotonic lysis buffer (10 mM HEPES pH7.9, 60 mM KCl, 1.5 mM MgCl2, 1 mM EDTA, 1 mM DTT, 0.075% NP-40, 1x protease/phosphatase inhibitor cocktails, Roche) and incubated for 10 minutes at 4°C with rotation. Nuclei were pelleted by centrifugation (1200 rpm, 4°C) for 10 minutes. The cytoplasm was collected from the supernatant, re-centrifuged (13500 rpm, 4°C, 10 minutes) and the supernatant was collected as soluble cytoplasmic fraction. Nuclei were washed five times in 800 µl hypotonic lysis buffer without NP-40 and lysed in 1 volume of nuclear lysis buffer (20 mM HEPES pH7.9, 400 mM NaCl, 1.5 mM MgCl2, 0.2 mM EDTA, 1 mM DTT, 5% glycerol, 1x protease/phosphatase inhibitor cocktails, Roche). Nuclear lysates were diluted with 2 volumes of dilution buffer (20 mM HEPES pH7.9, 1.6% triton x-100, 0.2% sodium deoxycholate, 1x protease/phosphatase inhibitor cocktails, Roche), followed by 10 sec sonication with a Bioruptor (Diagenode) at low energy and incubation with 10 U Benzonase (Sigma) for 5 min. Lysates were centrifuged (13500 rpm, 4°C, 10 minutes) and the supernatant was collected as soluble nuclear fraction. 50% of subcellular fractions were boiled in 1 volume of sample buffer at 95°C for 5 minutes, sonicated and analysed by immunoblotting. 50% of subcellular fractions was subjected to TRIzol extraction as above. The RNA from fractions was assessed qualitatively on a Fragment Analyzer (Advanced Analytical).

### Sucrose gradients

For preparation of sucrose gradients, 6 mL of 5% sucrose solution (5% sucrose in 10 mM tris-HCl pH7.5, 1 mM EDTA, 100 mM NaCl) was pipetted in an ultracentrifuge tube (Beckman). Then 6 mL of 50% sucrose solution (50% sucrose in 10 mM tris-HCl pH 7.5, 1 mM EDTA, 100 mM NaCl) were layered to the bottom of the tube by releasing the solution from a syringe fitted with a long blunt-end needle. The gradient was mixed by gradient maker (Gradient Master 108, Biocomp, program sucrose 5-50%) and kept at 4°C until usage. To prepare whole cell lysates, cells were trypsinized, washed in PBS and counted in order to achieve the same number of cells among the groups (5x 10^7^ cells per condition) and then centrifuged (1200 rpm, 5 min). The pellet was resuspended in 100 µL lysis buffer (25 mM Tris-HCl pH7.4, 105 mM KCl, 0.5% NP-40, 2 mM EDTA, 1 mM NaF, 0.5 mM DTT, 100U Ribolock inhibitor, Thermo, 1x protease/phosphatase inhibitor, Roche) per 2x 10^7^ cells. Lysates were incubated 30 min on ice with vortexing every 5 min, flash-frozen in liquid nitrogen, thawed, homogenized three times with a needle (BD Microlane 3, No. 14, 0.6 x 30 mm, blue), centrifuged (13000 rpm, 10 min, 4°C) and transferred to fresh tubes. RNase treatment was performed by incubating samples with 2.5 µL RNase A (10 mg/mL, Thermo), 2.5 µL RNase I (10 U/µL, Thermo), 2.5µL RNase T1 (1000 U/µL, Thermo), 5 µL RNase H (5 U/µL, NEB), 2.5 µL RNase III (1 U/µL, Thermo) for 1h at 4°C with rotation. 5% of supernatant was kept as input and resuspended with 1 volume of sample buffer, the rest was loaded on top of sucrose gradient and ultracentrifuged (30000 rpm, 4°C, 18h, stop DECEL, slow, SW40/1Ti swinging bucket rotor, Optime L-90K, Beckman). 24 fractions were taken from top to bottom of the tube as 500 µL aliquots and precipitated by incubation with 50 µL of 0.15% sodium deoxycholate for 10 min at RT, then mixed with 25 µl of 100% trichloroacetic acid and incubated 30 min, 4°C. Peptides were pelleted (12000 rpm, 15 min, 4°C), washed with 500 µL ice-cold acetone and pelleted again. Pellets were air-dried, resuspended in 40 µL sample buffer and analysed by immunoblotting. To assess RNA from sucrose gradients, samples from pooled fractions or inputs were subjected to TRIzol extraction and the recovered RNA was quantified by RT-qPCR.

### RNA mass spectrometry (RNA-MS)

Total RNA (5 µg in 20 µL ddH2O) was incubated with 2µL 100U Nuclease P1 (NEB) and 2 µL 10x Nuclease P1 buffer (NEB) for 30 min at 37°C, followed by inactivation of the enzyme (10 min, 75°C). For dephosphorylation of digested RNA, 1 µL 1U recombinant shrimp alkaline phosphatase (NEB) and 2.5 µL 10x CutSmart buffer (NEB) were added and incubated for 30 min at 37°C, followed by inactivation of the enzyme (5 min, 65°C). To assess methyladenosines in fluid samples, 20 µl of hydrolyzed and dephosphorylated RNA sample were diluted with 500 µl MeOH/H2O (80/20) and transferred to activated (by flushing with 0.5 ml CH3CN) and equilibrated (by flushing with 0.5 ml MeOH/H2O (80/20, v/v)) SPE column (Strata C18-E, 50 mg / 1 ml, Phenomenex). After passing the column, residual metabolites were eluted with 180 µl MeOH/H2O (80/20, v/v). The combined eluates were taken to dryness in a centrifugal evaporator. Samples were reconstituted in 50 µl of MeOH/H2O/AcOH (50/50/0.1, v/v/v). LC/MS analysis was performed on a Dionex Ultimate 3000 UHPLC system connected to a Q Exactive mass spectrometer (QE-MS) equipped with a HESI probe (Thermo). Chromatographic separation was achieved by applying 3 µl sample on a Hypercarb column (50 × 2.1 mm, 2.2 µm) (Thermo), protected by a Supelco ColumnSaver particle filter (Merck) and a gradient of mobile phase A (5 mM NH4OAc in CH3CN/H2O (40/60, v/v)) and mobile phase B (5 mM NH4OAc in CH3CN/H2O (495/5, v/v)) maintaining a flow rate of 200 µl/min and a column temp. of 45 °C. The LC gradient program was 5% mobile phase B for 2 min, followed by a linear increase to 100% B within 23 min, maintaining 100% B for 5 min and returning to 5% B in 2 min, followed by 7 min 5% B for column equilibration before each injection. The eluent was directed to the QE-MS from 7 min to 20 min after sample application. Mass detection was conducted in full scan pos. mode (at 70k resol., scan range m/z 220 - 320, AGC target 1E6 and 200 ms max. injection time). HESI parameters: Sheath gas: 20, aux gas: 1, spray voltage: 3.0 kV, capillary temp.: 300 °C, S-lens RF level: 50.0, aux gas heater temp.: 120 °C. Manual curation and integration of chromatographic peaks were performed with TraceFinder 5.1 using a mass tolerance of +/- 2 mMUs.

### Chromatin immunoprecipitation (ChIP) and CUT&RUN-sequencing

For ChIP, cells were fixed with 1% formaldehyde (10 min, 37°C), quenched in 0.125 M glycine (10 min, 37°C), washed in PBS and centrifuged (2000 rpm, 5 min). Pellets were resuspended in 500 µL cold cell lysis buffer (5 mM PIPES pH8.0, 85 mM KCl, 0.5% NP-40, 1x protease/phosphatase inhibitor, Roche) and lysed (10 min on ice). Nuclei were centrifuged (3000 rpm, 5 min) and resuspended in 300 µL cold nuclear lysis buffer (1% SDS, 10 mM EDTA, 50 mM tris-HCl pH8.0, 1x protease/phosphatase inhibitor, Roche) and lysed (10 min on ice). Lysates were sonicated (5 times 5 min, 30 sec on/off) with a Bioruptor (Diagenode) and pelleted (13000 rpm, 10 min). The supernatant was mixed with 2 mL dilution buffer (0.01% SDS, 1.1% triton x-100, 1.2 mM EDTA, 16.7 mM tris-HCl pH8.0, 167 mM NaCl, 1x protease/phosphatase inhibitor, Roche). Diluted samples were aliquoted, 5 µg antibodies were added (IP sample) or not (input) and incubated overnight (4°C with rotation). For pull-down, 20 µL of protein G dynabeads were added to IP samples, incubated (1.5 h with rotation), immobilized on a magnet and washed with wash buffer A (0.1% SDS, 1% triton x-100, 2 mM EDTA, 20 mM tris-HCl pH8.0, 150 mM NaCl), B (0.1% SDS, 1% triton x-100, 2 mM EDTA,20 mM tris-HCl pH8.0, 500 mM NaCl), C (0.25 M LiCl, 1% NP-40, 1% sodium deoxycholate, 1 mM EDTA and 10 mM tris-HCl pH8.0), and twice with D (10 mM tris-HCl pH8.0, 1 mM EDTA). For elution, samples were incubated with 500 µL elution buffer (1% SDS, 0.1 M NaHCO3) for 30 min with rotation. Reversal of cross-links was performed at 65°C overnight after adding 30 µL 5 M NaCl, 1 µL 10 mg/mL RNase A (Sigma), 10 µL 0.5 M EDTA, 20 µL 1 M tris-HCl pH6.8, 2 µL 10 mg/mL proteinase K (Sigma) to input and IP samples. DNA was purified by phenol/chloroform extraction, recovered in ddH2O and assessed by qPCR with selective primers (Table S5). For CUT&RUN-seq cells were harvested with accutase (Sigma), centrifuged (2500 rpm, 3 min) and washed three times in 1.5 mL wash buffer (20 mM HEPES pH7.5, 150 mM NaCl, 0.5 mM spermidine). Cells were incubated (10 min, RT) with 10 µL concanavalinA-coated magnetic beads (BioMag) resuspended in 1 volume of binding buffer (20 mM HEPES pH7.5, 10 mM KCl, 1 mM CaCl2, 1 mM MnCl2), immobilized on a magnet, permeabilized with 150 µL antibody buffer (20 mM HEPES pH7.5, 150 mM NaCl, 0.5 mM spermidine, 0.05% digitonin, 2 mM EDTA) and incubated with 1 µg primary antibody (800 rpm, 4°C, overnight with rotation). Samples were placed on a magnet, washed two times with 1 mL dig-wash buffer (20 mM HEPES pH7.5, 150 mM NaCl, 0.5 mM spermidine, 0.05% digitonin) and incubated (1 h, 800 rpm, 4°C with rotation) with 150 µL protein A/G-micrococcal nuclease (MNase) fusion protein (1 µg/mL, CST). Reactions were placed on a magnet, washed two times with 1 mL dig-wash buffer and once with 1 mL rinse buffer (20 mM HEPES pH7.5, 0.05% digitonin, 0.5 mM spermidine). For chromatin digestion and release, samples were incubated (30 min, on ice) in ice-cold digestion buffer (3.5 mM HEPES pH7.5, 10 mM CaCl2, 0.05% digitonin). The reaction was stopped by addition of 200 µL stop buffer (170 mM NaCl, 20 mM EGTA, 0.05% digitonin, 50 µg/mL RNaseA, 25 µg/mL glycogen) and fragments were released by incubation (30 min, 37°C). The supernatant was incubated (1 h, 50°C) with 2 µL 10% SDS and 5 µL proteinase K (10 mg/mL, Sigma). Chromatin was recovered by phenol/chloroform extraction and resuspended in 30 µL TE (1 mM tris-HCl pH8.0, 0.1 mM EDTA). For sequencing, biological replicates were quantified with a Fragment Analyzer (Advanced Analytical), pooled and subjected to library preparation. Libraries for small DNA fragments (25-75 bp) were prepared based on the NEBNext Ultra II DNA library prep Kit for Illumina (NEB#E7645).

### Capture hybridization analysis of RNA targets (CHART) and CHART-seq

Cells were fixed with 1% formaldehyde (6 min, 37°C), quenched in 0.125 M glycine (10 min, 37°C), washed in PBS and centrifuged (2000 rpm, 5 min). Pellets were resuspended in 300 µL cold cell lysis buffer (5 mM PIPES pH8.0, 85 mM KCl, 0.5% NP-40, 100U Ribolock inhibitor, Thermo, 1x protease/phosphatase inhibitor, Roche) and lysed (10 min on ice). Nuclei were centrifuged (3000 rpm, 5 min) and resuspended in 50 µL cold nuclear lysis buffer (1% SDS, 10 mM EDTA, 50 mM tris-HCl pH8.0, 100U Ribolock inhibitor, Thermo, 1x protease/phosphatase inhibitor, Roche) and lysed (10 min, RT). Lysates were sonicated (5 times 5 min, 30 sec on/off) with a Bioruptor (Diagenode) and pelleted (13000 rpm, 10 min). 10% of supernatant was kept as input and stored at -20°C. The remaining supernatant was diluted in 1 mL pre-warmed capturing buffer (0.5 M LiCl, 4 M urea, 100U Ribolock inhibitor, Thermo, 1x protease/phosphatase inhibitor, Roche) and 10 µL from an equimolar stock (10 µM) of six biotin-tagged capturing oligonucleotides (Table S6) were added or not. Following incubation of lysates with 100 nM capturing oligos for 3h at 65°C with shaking, 30 µL streptavidin C1 dynabeads (Thermo) were equilibrated and added to each sample for pull-down (45 min, RT with rotation). For elution, samples were incubated with 500 µL elution buffer (1% SDS, 0.1 M NaHCO3) for 30 min with rotation. Reversal of cross-links was performed at 65°C overnight after adding 30 µL 5 M NaCl, 1 µL 10 mg/mL RNase A (Sigma), 10 µL 0.5 M EDTA, 20 µL 1 M tris-HCl pH6.8, 2 µL 10 mg/mL proteinase K (Sigma) to all samples. Chromatin was purified by phenol/chloroform extraction, resuspended in ddH2O and assessed by qPCR. To prepare samples for sequencing, purified chromatin was resuspended in 30 µL TE (1 mM tris-HCl pH8.0, 0.1 mM EDTA). Samples were quantified with a Fragment Analyzer (Advanced Analytical) and pooled biological replicates were subjected to library preparation. Libraries for DNA fragments were prepared based on the NEBNext Ultra II DNA library prep Kit for Illumina (NEB#E7645).

### Generation of FASTQ, BAM and bedgraph files

For CUT&RUN-seq, base calling was performed using Illumina’s FASTQ Generation software v1.0.0 and sequencing quality was tested using FastQC. Reads were mapped with Bowtie2 v2.3.5.1(47) to human hg19, human T2T or mouse mm10 genome. CUT&RUN-seq samples were normalized to the number of mapped reads in the samples with the least number of reads. BAM files obtained after read-normalization were sorted and indexed using SAMtools v1.9 and Bedgraph files were generated using the genomecov function from BEDTools v2.26.0(48). The Integrated Genome Browser (IGB) was used to visualize density files.

### Generation of metagene plots, heatmap and bar charts

CUT&RUN-seq metagene plots were generated using the R package ‘metagene’ with the assay parameters ‘ChIPseq’ and 100 bins, testing 80 accessible and 1123 predicted AsiSI sites. Read count was performed using BEDTools. Data from the heatmap and read count analyses were visualized with RStudio (v2022.7.2.576) using package ‘ggplot2’.

### Single-end enhanced cross-linking immunoprecipitation (seCLIP)-sequencing

For crosslinking of ribonucleoprotein complexes in vivo, cells were washed in PBS, aspirated, irradiated by UV-light (254 nm, 400 mJ/cm^2^), resuspended in ice-cold PBS, scraped and centrifuged (1200 rpm, 5 min). The pellets were stored at -80°C. For lysis, the cells were incubated in 1 mL seCLIP lysis buffer (50 mM tris-HCl pH7.4, 100 mM NaCl, 1% NP-40, 0.1% SDS, 0.5% sodium deoxycholate, 440U murine RNase inhibitor, Thermo, 1x protease/phosphatase inhibitor, Roche) for 5 min at 4°C and lysates were sonicated (1x 5 min, 30 sec on/off) with a Bioruptor (Diagenode) followed by digestion with 40U RNaseI (Invitrogen) and 10U TURBO DNase (Invitrogen) for 5 min at 37°C with rotation and centrifugation (13000 rpm, 10 min). The supernatant was collected for immunoselection, the remaining supernatant was incubated with 15 µg antibodies, which were pre-conjugated to 1:1 mix of 63 µL protein A/G dynabeads (Invitrogen), and incubated (4°C, overnight with rotation). The immunocomplexes were captured on a magnet, was washed twice in 800 µL high-salt wash buffer (50 mM tris-HCl pH7.4, 1 M NaCl, 1% NP-40, 1 mM EDTA, 0.1% SDS, 0.5% sodium deoxycholate), followed by two washes in 800 µL wash buffer (20 mM tris-HCl pH7.4, 10 mM MgCl2, 0.2% tween-20, 5 mM NaCl). Subsequent library preparation was performed as described(49).

### Bioinformatic analysis of seCLIP-seq

seCLIP samples were processed based on published eCLIP analysis protocols (49, 50). Briefly, adapters were trimmed twice using cutadapt (v1.14). Trimmed FASTQ files were then first aligned to a genome index consisting only of Repbase annotated repetitive elements using the STAR aligner (v2.7.6a). Reads that did not map to repetitive elements were then aligned to hg19 using STAR. Genome mapped BAM files were sorted with Samtools and PCR duplicates were removed by a custom python script(49, 50). The read counts of all hg19 aligned bam files were counted and the sample with the lowest count was used to normalize all bam files. Sequencing depth-normalized bam files were then converted to bigwig files with deeptools (v3.5.1) bamCoverage. Bigwig files of seCLIP-seq biological replicates were merged by calculating the average between the two biological replicates using deeptools bigwigCompare (--scaleFactors 1:1 --operation mean). Log2 fold-change bigwig files between merged seCLIP-seq replicates and corresponding inputs were generated with deeptools bigwigCompare (-- scaleFactors 1:1 --operation log2 --pseudocount 1).

### Dimethyl sulfate mutational profiling combined with Nanopore sequencing (Nano-DMS-MaP-seq)

For *in vivo* structural probing of RNA, cells were incubated with 25 mM DMS for 6 min at 37°C, placed on ice, washed with cold PBS containing 1% β-mercaptoethanol, harvested by scraping, centrifuged (1200 rpm, 5 min) and resuspended in TRIzol to recover RNA. After RNA extraction, 10 µg of total RNA was treated with Turbo DNase (Invitrogen) and subsequently purified using NucleoSpin PCR Clean-up columns (Macherey Nagel). NEAT1 cDNA was synthesized with the Marathon RT probing protocol as described(51) using an anchored polydT primer (TTTTTTTTTTTTTTTTTTTTVN). cDNA was amplified by PCR with selective primers (PCR A: fwd, 5’-GGAGTTAGCGACAGGGAGGGATG-3’; rev, 5’-AGAACAAAAGAGCACTACCGGTGTAC-3’; PCR B: fwd, 5’-ATTTGTGCTGTAAAGGGGAAGAAAAGTGATTAG-3’; rev, 5’-

TCTGTGTGTGAGAAATGGCAGGTCTAG-3’). Subsequent library preparation and analysis steps were performed as described(51), with sequencing being carried out with the ONT native barcoding kit 14 (SQK-NBD114.96) on a MinION R10.4.1 flow cell.

## RESULTS

### DNA damage increases the levels of NEAT1 isoforms

NEAT1 transcripts are upregulated in about 65% of all tumours(36, 37, 52). We hypothesized that high levels of NEAT1 protect tumour cells from excessive DNA damage and used RT-qPCR to quantify the total amount of NEAT1 (NEAT1_1+NEAT1_2, hereinafter NEAT1) and the level of the NEAT1_2 isoform in human U2OS cells the absence or presence of etoposide. Incubation with etoposide elevated both NEAT1 and NEAT1_2 levels ∼5-fold in the nuclear, but not the cytoplasmic fraction (Fig. 1A; Fig. S1A). The onset of DNA damage was confirmed by immunoblotting for the DNA damage marker ser-139 phosphorylated histone H2.X variant (γH2A.X) (Fig. S1B). The etoposide-responsive NEAT1 induction phenotype could also be observed in HEK293 cells (Fig. S1C). NEAT1 levels were further assessed by RNA-FISH and were increased upon treatment with etoposide (Fig. 1B). The integrity of the RNA-FISH assay was confirmed by colocalization of NEAT1-probes with the *bona fide* NEAT1 interactor SFPQ (Fig. S1D). Thus, etoposide treatment elevates NEAT1 levels in human cells.

**Figure 1.**
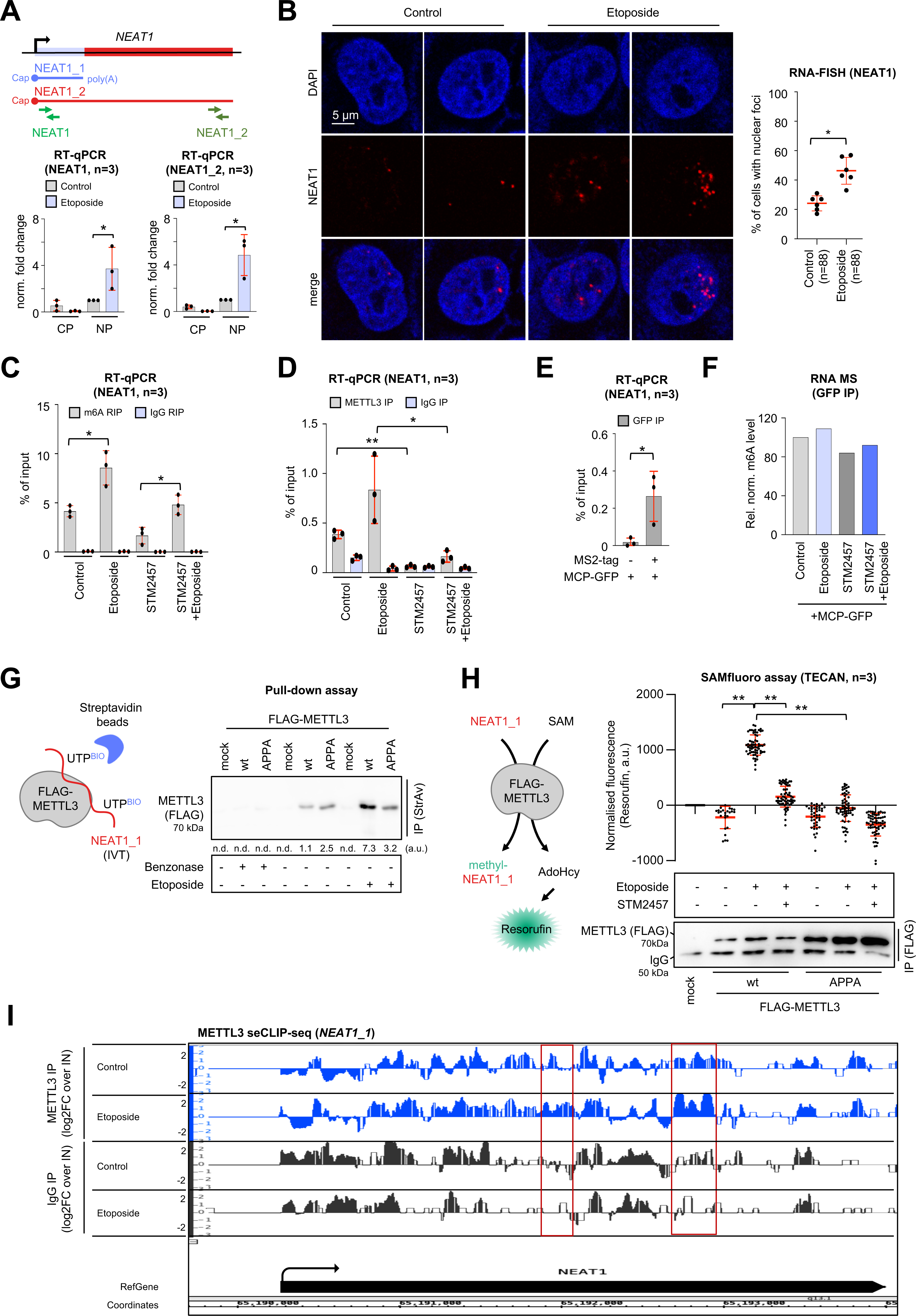
DNA damage elevates the levels and METTL3-dependent methylation of NEAT1 transcripts. (**A**) Scheme (top) and RT-qPCR (bottom) assessing transcript levels of NEAT1 isoforms from RNA upon subcellular fractionation into cytoplasm (CP) and nucleoplasm (NP) of U2OS cells. Green arrowhead, RT-qPCR primer. (**B**) Imaging (left) and quantitation (right) of Quasar570-labelled RNA-FISH probe signals in U2OS cells. Each dot represents an average from two acquisitions. (**C, D**) RT-qPCR assessing NEAT1 levels upon m6A RNA immunoprecipitation (RIP) from total RNA (C) or METTL3 immunoprecipitation (IP) from whole cell lysates (D) of U2OS cells with selective antibodies. IgG, immunoglobulin G control. (**E**) RT-qPCR assessing NEAT1 levels upon ectopic expression of GFP-tagged MS2 coat protein (MCP-GFP) and immunoselection from whole cell lysates of wild type HEK293 (-MS2-tag) or HEK293:24xMS2-NEAT1 (+MS2-tag) cells. (**F**) Quantitation of N6-methyladenosine (m6A) levels by RNA mass spectrometry upon ectopic expression of MCP-GFP and immunoselection from whole cell lysates of HEK293:24xMS2-NEAT1 cells. (**G**) Scheme of pull-down assay (left) and immunoblot (right) displaying ectopically expressed FLAG-tagged METTL3 variants upon immunoprecipitation with biotin (BIO)-labelled and immobilized NEAT1_1 *in vitro* transcription (IVT) product. a.u., arbitrary units; n.d., not detectable. (**H**) Scheme of SAMfluoro assay (left) and fluorescent counts (right) displaying Resorufin levels. Immunoblot (bottom), loading control; AdoHcy, S-adenosine-L-homocysteine; mock, non-transfected lysate. (**I**) Browser tracks of METTL3 seCLIP-seq reads at the *NEAT1_1* locus in U2OS cells. Read log2 fold-changes between merged seCLIP-seq duplicates and size-matched input are shown. Red box, region of interest; arrowhead, transcription start site. */**, p-value <0.05/ <0.001; two-tailed t-test. Error bar, mean ±SD. Representative images are shown. n=number of biological replicates or imaged cells.

### METTL3 places m6A marks on NEAT1 upon DNA damage

The N6-methyladenosine (m6A) RNA modification is rapidly induced by UV-irradiation(53). Interestingly, four m6A sites have been mapped to the 5’ end region NEAT1 that overlaps between NEAT1_and NEAT1_2(54). The m6A marks are placed by the DSB-inducible RNA methyltransferase 3 (METTL3), a direct target of ATM, and stabilise NEAT1(55, 56). We speculated that METTL3 places m6A marks on NEAT1 upon DNA damage. First, we wished to test if the DDR indeed engages METTL3 in U2OS cells. METTL3 accumulates at DSBs upon phosphorylation(55). Lacking a phospho-specific METTL3 antibody, we tested for colocalization of METTL3 with DSB foci as a proxy for METTL3 activation. We incubated cells in the absence or presence of etoposide, or preincubated them with the selective ATM inhibitor KU-55933 and assessed the subcellular localization of METTL3 and γH2A.X by PLAs and confocal microscopy. Indeed, we measured a strong increase in METTL3/γH2A.X PLA signals upon etoposide treatment, which was sensitive to the ATM inhibitor and suggests DNA damage-induced accumulation of METTL3 at DSBs (Fig. S1E). Partial colocalization of METTL3/γH2A.X antibody signals upon etoposide treatment was also visible by indirect immunofluorescence (Fig. S1F). We also performed dot blots with a selective m6A antibody and observed >5-fold increase in m6A antibody reactivity on total RNA from etoposide-treated cells (Fig. S1G). This suggests that METTL3 is responsive to etoposide incubation in U2OS cells. To assess m6A marks on NEAT1, we incubated U2OS cells in the absence or presence of etoposide, or pre-treated them with the validated METTL3 inhibitor STM2457(57) and performed RIP-RT-qPCR with the m6A antibody (Fig. 1C). The m6A antibody selectively enriched NEAT1 in all 4 conditions. Strikingly, however, NEAT1 enrichment was particularly strong upon etoposide treatment and partially sensitive to STM2457 preincubation. Importantly, the RNA integrity was monitored by urea-PAGE (Fig. S2A). Next, we employed a selective METTL3 antibody for immunoprecipitation and asked if NEAT1 is differentially associated with METTL3 upon DNA damage. Again, we found significant amounts of NEAT1 coenriched with METTL3, which were modestly increased by etoposide incubation and sensitive to STM2457 pre-treatment (Fig. 1D; Fig. S2B). As expected, pre-treatment with STM2457 prevented the etoposide-responsive induction of NEAT1 levels, but had little impact on the NEAT1 steady-state levels in unperturbed cells (Fig S2C). We wished to corroborate our findings in a different system and employed CRISPR/Cas9 technology to introduce an array of 24 stem loop-forming MS2 RNA-binding sequences (MS2-tag) upstream of the TSS of the *NEAT1* locus in HEK293 cells. The heterozygous knock-in of the MS2-tag was validated by PCR in a subset of clones and generated the monoclonal cell line HEK293:24xMS2-NEAT1 (Fig. S2D). To test if MS2-NEAT1 transcripts could be enriched from HEK293:24xMS2-NEAT1 cells, we expressed GFP-tagged MS2 coat protein (MCP-GFP) and used a GFP antibody for immunoselection (Fig. S2E). Indeed, the selective enrichment of NEAT1 from HEK293:24xMS2-NEAT1, but not wild type HEK293 cells was confirmed by RT-qPCR (Fig. 1E). Next, we used RNA mass spectrometry to determine the relative abundance of m6A marks on immunoselected MS2-NEAT1 (Fig. 1F). Reassuringly, m6A marks were modestly increased upon etoposide incubation and sensitive to STM2457 preincubation, further suggesting that METTL3 places m6A marks on NEAT1.

To test if METTL3 binds NEAT1 *in vitro*, we immobilized *in vitro* transcribed, UTP-biotinylated NEAT1_1 on streptavidin-coated beads and incubated the RNA with whole cell lysates from HEK293 cells expressing wild type FLAG-METTL3 or a catalytic inactive point mutant (APPA) in the absence or presence of etoposide. We found increased binding of wild type, but not APPA FLAG-METTL3 upon treatment with etoposide, which was sensitive to benzonase digestion, in streptavidin pull-downs (Fig. 1G). Next, we performed a fluorescence-based SAMfluoro *in vitro* methylation assay (Fig. 1H). We immunoselected wild type or APPA mutant FLAG-METTL3, which were again ectopically expressed in HEK293 cells in the absence or presence of etoposide or STM2457, from whole cell lysates and incubated the samples with *in vitro* transcribed, non-methylated NEAT1_1 and SAMfluoro reagents. The reaction product Resorufin was quantified as a proxy for methyltransferase activity. We found elevated levels of Resorufin upon immunoselection of wild type, but not mutant FLAG-METTL3 from etoposide treated cells, which were sensitive to STM2457. The immunoselection of FLAG-METTL3 variants and the integrity of *in vitro* transcribed NEAT1_1 were confirmed by immunoblotting and agarose gel electrophoresis, respectively (Figs. S2F, S2G). To test if METTL3 binding to NEAT1 is modulated by DNA damage *in vivo*, we performed seCLIP-seq and determined several regions within NEAT1_2 that displayed enrichment of reads upon METTL3 immunoselection irrespective of etoposide treatment (Fig. S2H). Moreover, METTL3 to NEAT1 seems to be selectively increased upon etoposide treatment at a subset of regions, including the 3’ end region of NEAT1_1 (Fig. 1I), further suggesting enhanced binding of METTL3 to NEAT1 upon DNA damage. We conclude that NEAT1 increases in levels and is hyper-methylated by METTL3 upon DNA damage.

### The depletion of NEAT1 impairs the DDR

METTL3 has recently been described as amplifier of the tumour suppressor protein p53-dependent stress response in mouse embryonic fibroblasts(58). Murine METTL3 stabilizes p53 to promote the transactivation of target genes and places m6A marks on a subset of p53-responsive transcripts, including *NEAT1* upon treatment with the DNA intercalator doxorubicin. To test if METTL3 and NEAT1 display genetic interaction upon DNA damage in human cells, we utilized the plasmid-based DSBR reporter construct DR-GFP, which contains an inactive *GFP* reporter locus and a target site for the restriction enzyme I-SceI to cleave the reporter *in vivo* and prevent GFP expression. The repair of the I-SceI restriction site restores functional GFP expression and can be used as a proxy for HR-mediated DSBR efficacy. As expected, we observed expression of the GFP reporter product upon cotransfection of DR-GFP- and I-SceI-encoding plasmids in HEK293 cells (Fig. S3A). In contrast, GFP expression was markedly reduced upon combining the transfection of reporter plasmids with a pool of NEAT1-selective siRNA or siRNA to deplete METTL3. Next, we combined etoposide treatment with the depletion of NEAT1 and METTL3 and assessed the amount of broken chromatin by neutral comet assays in U2OS cells (Fig. 2A). As expected, the depletion of NEAT1 and METTL3 did not cause the formation of broken chromatin in the absence of etoposide and the incubation with etoposide induced prominent formation of DNA tails. However, the depletion of both NEAT1 and METTL3 enhanced the etoposide-induced tail phenotype and could be observed most prominently upon co-depletion of NEAT1 and METTL3. The siRNA knockdown efficacy of NEAT1 and METTL3 was monitored by RT-qPCR and immunoblotting (Figs. S3B, S3C). This suggests that NEAT1 and METTL3 promote the efficient repair of etoposide-induced DSBs. Next, we performed etoposide pulse-chase kinetics and used confocal imaging to assess the impact of NEAT1 depletion on the formation and clearance of 53BP1-positive DSB foci (Fig. 2B). Surprisingly, the depletion of NEAT1 by siRNA impaired the formation of such foci. Next, we depleted NEAT1 by DNA-RNA chimeric antisense oligonucleotides (ASOs) and assessed DSB signalling efficacy by immunoblotting (Fig. 2C; Fig. S3D). Indeed, the depletion of NEAT1 with pooled ASOs impaired the efficient phosphorylation of ATM/ATR substrate and the accumulation of γH2A.X marks in etoposide pulse-chase kinetics. Likewise, we observed defects in the formation of etoposide-induced γH2A.X-positive DSBs foci in NEAT1-deficient cells (Fig. 2D). We noticed that the transfection of NEAT1_2 targeting ASOs depleted both NEAT1_1 and NEAT1_2 to similar extent, which may perhaps reflect feedback inhibition of RNA synthesis at the *NEAT1* locus, but also raised concerns about viability and cell cycle-dependent secondary effects upon ASO transfection. To test the latter, we performed FACS analysis and compared the cell cycle profiles of NEAT1-proficient and -deficient U2OS cells (Fig. S3E). We found that the depletion of NEAT1 modestly increased the number of cells in G1-phase, but neither impaired progression through S-phase, nor triggered the appearance of cells in sub-G1-phase. We conclude that NEAT1 promotes efficient DSB signalling and genome stability.

**Figure 2.**
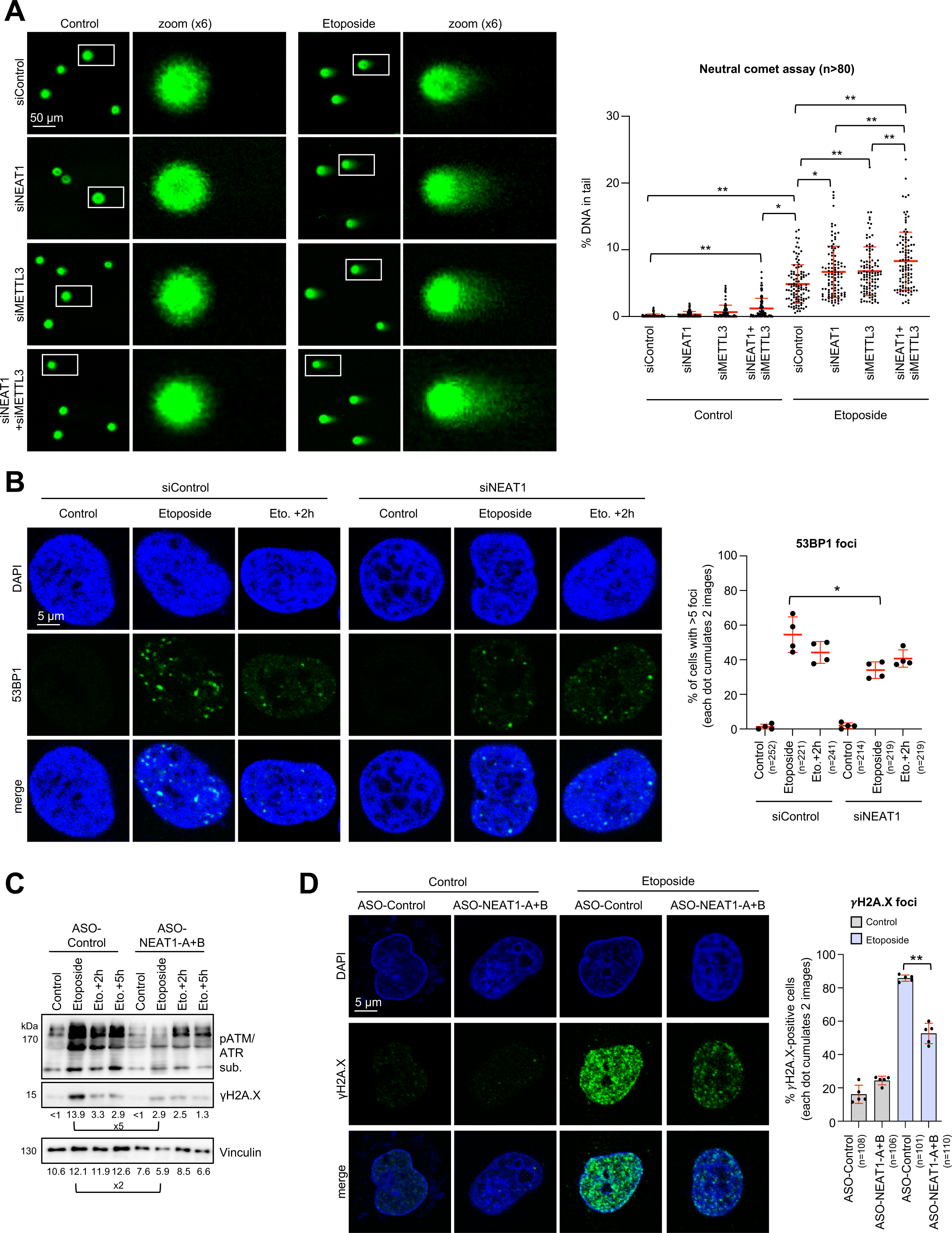
Depletion of NEAT1 elevates DNA damage and impairs the DDR in U2OS cells. (**A**) Imaging (left) and quantitation (right) of neutral comet assay displaying SYBR gold-stained DNA upon transfection of siRNA. White box, zoom. Each dot represents one measurement. (**B**) Imaging (left) and quantitation (right) of 53BP1 signals upon transfection of siRNA. Each dot represents an average from two acquisitions. (**C**) Immunoblots detecting phospho (p)ATM/ATR substrates and serine-139 phosphorylated H2A.X (γH2A.X) upon transfection of antisense oligonucleotides (ASOs). Vinculin, loading control. (**D**) Imaging (left) and quantitation (right) of γH2A.X signals upon transfection of ASOs. Each dot represents an average from two acquisitions. */**, p-value <0.05/ <0.001; two-tailed t-test. Error bar, mean ±SD. Representative images are shown. n=number of imaged cells.

### NEAT1 accumulates at DSBs in a METTL3-dependent manner

NEAT1 associates with several hundred protein-coding gene promoters to regulate RNAPII transcriptional activity in human tissue culture cells(59, 60). We reasoned that NEAT1 chromatin association may be altered by DNA damage. To investigate NEAT1 in context of locus-specific DSBs, we employed U2OS DIvA cells, which stably express the 4-hydroxytamoxifen(4-OHT)-inducible endonuclease AsiSI (Fig. S4A). AsiSI cleavage produces ∼80 mostly promoter-associated DSBs(61). First, we monitored the induction of AsiSI-induced DSBs by immunoblotting and confocal imaging (Figs. S4B, S4C). We observed increased γH2A.X level and prominent formation of partially colocalising γH2A.X- and 53BP1-positive foci in DIvA, but not wild type cells upon incubation with 4-OHT. However, a modest increase in γH2A.X levels and some DSB foci were also detectable in DIvA in the absence of 4-OHT. Subsequently, we compared DIvA cells with 4-OHT-incubated wild type U2OS cells and performed manual ChIP for γH2A.X at the AsiSI site DS1 (*CCBL2/RBMXL1* promoter) for further validation (Fig. S4D). As expected, 4-OHT incubation of DIvA, but not wild type U2OS cells elevated γH2A.X levels up to 2000 nts upstream of the DS1 cleavage site. Next, we wished to confirm that the depletion of NEAT1 impairs the formation of γH2A.X-positive DSB foci also in the DIvA system, and indeed observed diminished γH2A.X foci intensities upon incubation with 4-OHT (Fig. 3A). Intriguingly, NEAT1 depletion did not prevent the formation of NBS1-positive foci. NBS1 is an upstream component of the DSB-sensing machinery. This suggests that the induction of DSBs by AsiSI *per se* is not impaired by NEAT1 depletion. Likewise, we observed loss of γH2A.X occupancy around the DS1 site in NEAT1-deficient cells (Fig. 3B). In contrast, NEAT1 depletion increased the levels of histone H2B lys-120 acetylation (H2B120ac), a DSB signalling marker upstream of γH2A.X(61), close to DS1 (Fig. 3C). These findings encouraged us to perform CHART-seq(60), which assesses the chromatin occupancy of lncRNA upon immunoselection with hybridized biotinylated DNA oligonucleotides and qPCR-based quantification of coenriched DNA (Fig. 3D). Strikingly, we coenriched DNA from a subset of promoter-associated DSBs in the presence of 4-OHT, but not upon STM2457 preincubation from DIvA cells (Fig. 3E). The enrichment of DNA was dependent on a pool NEAT1-selective capturing oligos and could not be observed in wild type U2OS cells. CHART-seq data were confirmed by inspection of browser tracks of the promoter-associated AsiSI target site *RNF8* (Fig. 3F), the originating *NEAT1_1* locus (Fig. S4E) and by manual CHART at DS1 (Fig. S4F). To corroborate our biochemical data, we employed RNA-FISH for confocal imaging-based localization studies of NEAT1. Reassuringly, we observed elevated NEAT1 probe signals upon treatment with etoposide, which partially colocalized with 53BP1-positive DSB foci and were sensitive to STM2457 preincubation (Fig. 3G). Next, we used the RNA-PLA assay, which generates signals upon combining a selective antibody with a selective DNA oligonucleotide probe(62). Indeed, etoposide treatment selectively increased RNA-PLA signals when combining γH2A.X staining with a NEAT1-selective probe (Fig. 3H; Fig. S4G). We conclude that NEAT1 accumulates at DSBs to promote DSB signalling in a METTL3-dependent manner.

**Figure 3.**
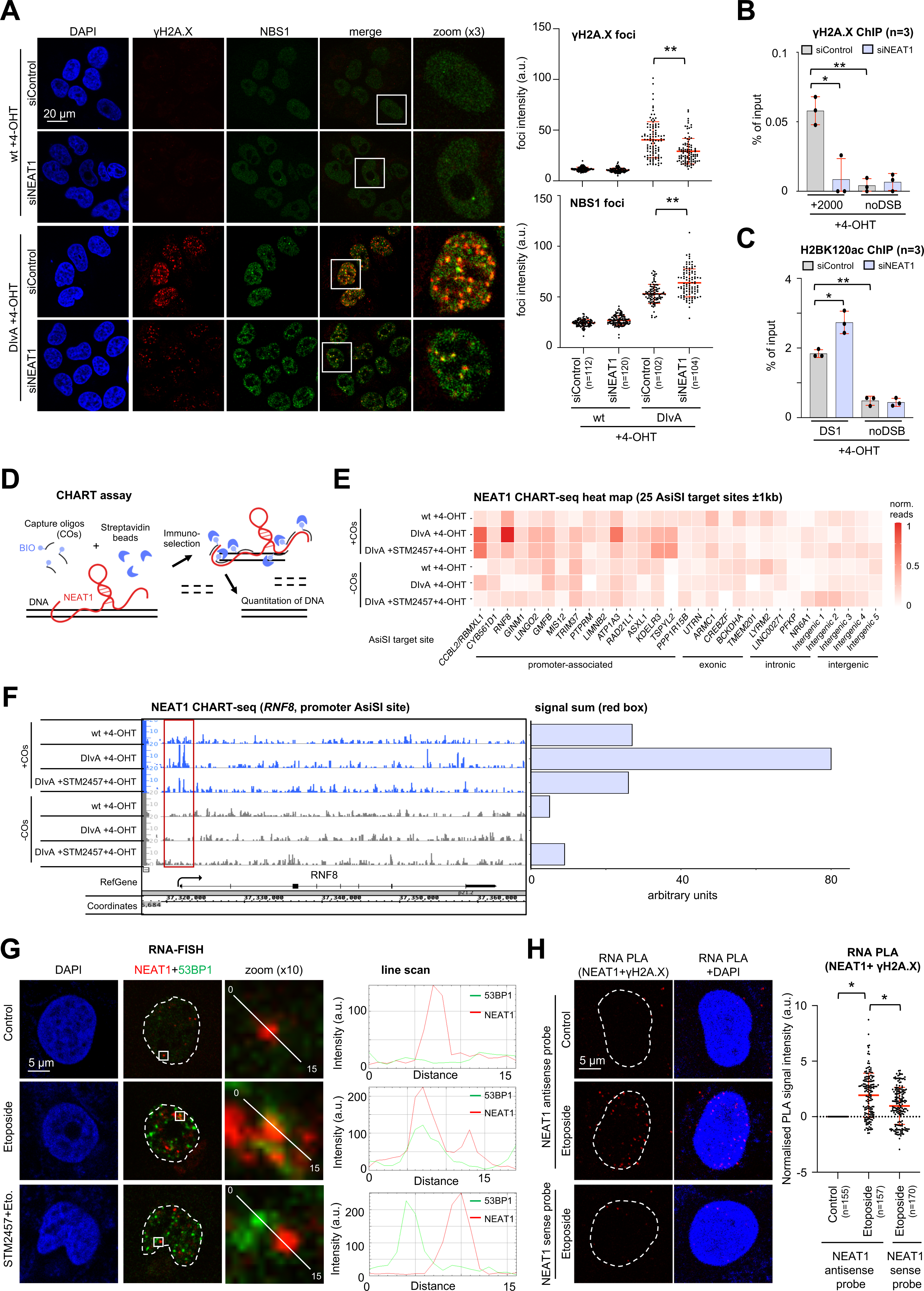
NEAT1 accumulates at DNA double-strand breaks to promote DDR signalling in U2OS cells. (**A**) Imaging (left) and quantitation (right) of γH2A.X and NBS1 signals in wild type (wt) and DIvA U2OS cells upon transfection of siRNA and incubation with 4-hydroxytamoxifen (4-OHT). White box, zoom. Each dot represents one measurement. (**B, C**) Manual ChIP for γH2A.X (B) and histone H2B lys-120 acetylation (H2BK120ac) (C) upon transfection of siRNA. +2000, distance from AsiSI target site in bps, noDSB, non-restricted control. (**D**) Scheme of CHART assay. BIO, biotin-tag (**E**) CHART-seq heatmap for read counts at 25 AsiSI target sites in wild type U2OS or DlvA cells. COs, capturing oligos. (**F**) NEAT1 CHART-seq browser tracks of *RNF8* gene. Red box, region of interest; arrowhead, transcription start site. (**G**) Imaging (left) and line scan quantitation of colocalization (right) of Quasar570-labelled RNA-FISH probe and 53BP1 antibody merged signals. Broken white circle, nucleus; white box, zoom. More than 80 cells were assessed. (**H**) Imaging (left) and quantitation (right) RNA-PLA signals. Each dot represents one measurement. Broken white circle, nucleus. */**, p-value <0.05/ <0.001; two-tailed t-test. Error bar, mean ±SD. Representative images are shown. a.u., arbitrary units; n=number of biological replicates or imaged cells.

### DNA damage alters NEAT1 secondary structure in a METTL3-dependent manner

The m6A modification is associated with alterations in the structure and function of lncRNA. m6A methylation of the metastasis-associated lung adenocarcinoma transcript 1 (MALAT1), for instance, triggers conformational changes that alter the structure of MALAT1 itself as well as the composition and function of MALAT1-dependent nuclear speckles(63, 64). To assess, if DNA damage alters the structure of NEAT1, we applied dimethyl sulfate mutational profiling combined with Nanopore sequencing (Nano-DMS-MaP-seq), which we have recently established in HEK293 cells(51). Nano-DMS-MaP-seq is an extension of the previously described DMS-MaP-seq method and allows *in vivo* RNA structural probing with long read sequencing(65). It relies on selective methylation of unpaired adenine and cytosine bases at their Watson-Crick face when they are located within accessible regions of RNA, such as in loops or single-stranded motifs. These methylations can trigger nucleotide exchanges upon reverse transcription that are mapped by Nanopore sequencing (Fig. 4A). Prior to sequencing, we tested primer binding sites to efficiently reverse transcribe NEAT1 from unprobed (-DMS) or DMS-probed (+DMS) HEK293 cells that were incubated in the absence or presence of etoposide or pre-treated with STM2457. Using two binding sites in the middle and 3’ end of NEAT1_1, we were able to generate cDNA that covers full length NEAT1_1 from all conditions (Fig. S5A). However, we failed to amplify cDNA for the NEAT1_2 isoform, likely due to the length, high GC-content, or abundant G-quadruplex structures within NEAT1_2(66). Next, we performed Nano-DMS-MaP-seq from two biological replicates. As expected, DMS probing was selectively modifying A and C residues, independently of etoposide or STM2457 pre-treatments (Fig. S5B). Quantitation and computational analysis of DMS reactivities revealed that NEAT1 contains a flexible region at the 3’ end that shows distinct folding upon distinct treatments (Fig. 4B). Importantly, the predicted RNA-RNA interactions in that region were selectively altered by etoposide and suppressed by preincubation with STM2457. The calculated Pearson correlation of DMS reactivities was close to 0.9 across all replicates (Fig. S5C), indicating high-quality structural probing data and supporting experimental reproducibility. We conclude that etoposide treatment alters the secondary structure at the 3’ end region of NEAT1_1 in a METTL3-dependent manner.

**Figure 4.**
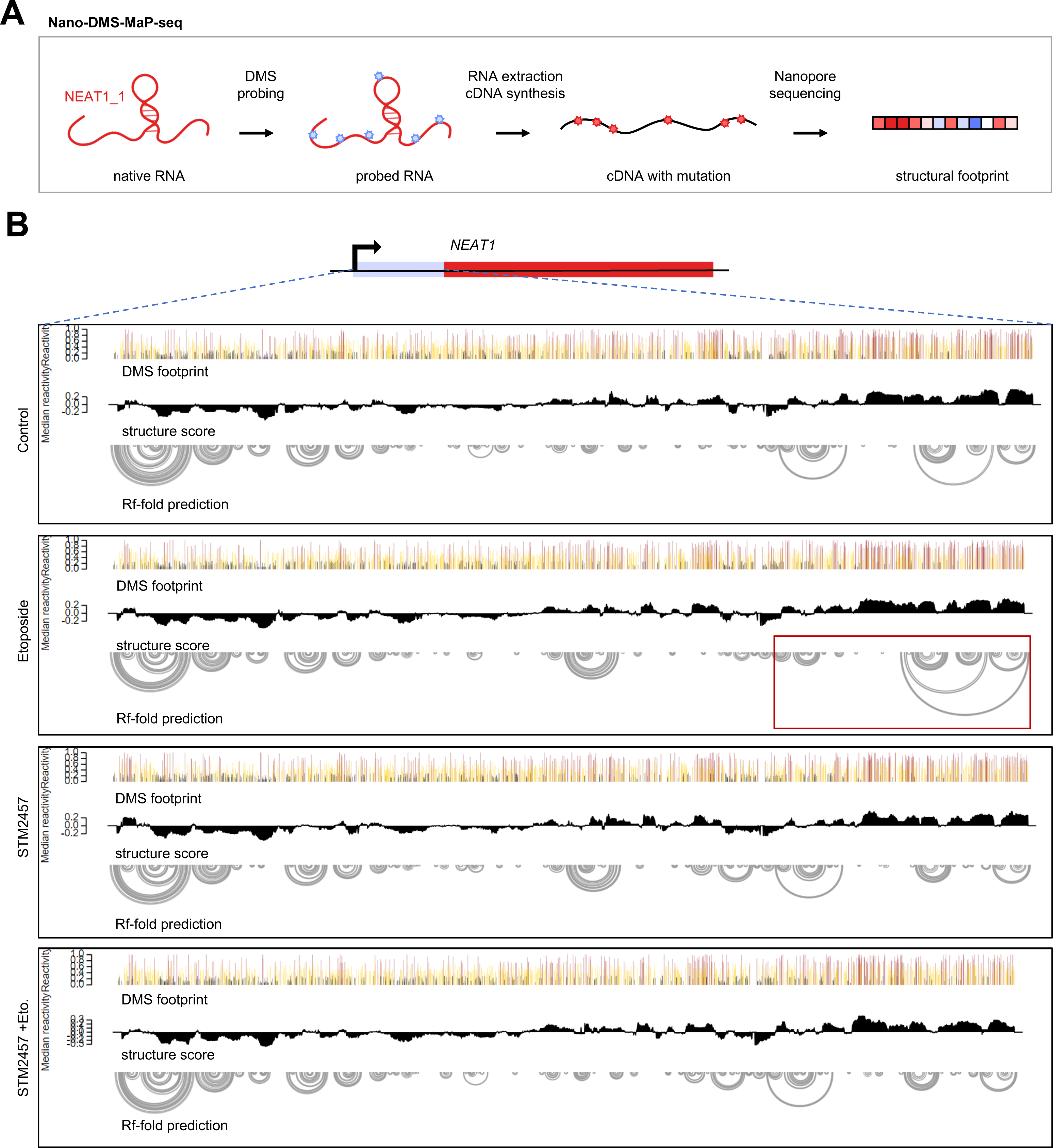
DNA damage alters NEAT1 secondary structure in HEK293 cells. (**A, B**) Scheme (A) and result (B) of the Nano-DMS-MaP-seq approach. Blue asterisk, DMS (dimethyl sulfate) modification; red asterisk, base conversion; red box, etoposide-responsive area.

### CHD4 associates with NEAT1 *in vivo* and *in vitro*

So far, our data suggest that METTL3 places m6A marks on NEAT1 to promote the accumulation of NEAT1 at promoter-associated DSBs, which likely is accompanied with alterations in NEAT1 secondary structure. We hypothesized that NEAT1 hyper-methylation may alter the association to RBPs that are involved in the DDR. To identify potential candidates, we merged a recently published NEAT1 interactome with a DNA-RNA hybrid (R-loop) interactome, as both NEAT1 and R-loop tend to accumulate close to promoter regions(67–69). Among the 64 identified candidates were the *bona fide* NEAT1 interactors NONO, SFPQ and PSPC1 as well as several RNA metabolic RBPs and components of the RNAPII machinery (Fig. 5A). Interestingly, the list of candidates also contained two DSBR factors (RAD50 and CHD4). The histone deacetylase CHD4 caught our attention as it is both validated as DSBR factor and binder of several lncRNA(70–72), including NEAT1(73). Using our HEK293:24xMS2-NEAT1 cells, we confirmed the selective enrichment of CHD4 with MCP-GFP by immunoblotting (Fig. 5B). To test if CHD4 interacts with NEAT1 directly, we incubated radio-labelled *in vitro* transcribed NEAT1_1 with CHD4, which was immunoselected from HEK293 whole cell lysates, and determined the amount of coimmunoprecipitated NEAT1_1 by autoradiography (Fig. 5C). Indeed, the co-enrichment of NEAT1_1 was dependent on CHD4 and sensitive to benzonase digestion. Next, we incubated equal amounts of whole cell lysates from HEK293 cells expressing recombinant, GFP-tagged full-length CHD4 (CHD4-GFP) with streptavidin-coated beads that were conjugated with or without *in vitro* transcribed, UTP-biotinylated NEAT1_1, and compared the amount of pulled down CDH4-GFP (Fig. 5D). Again, CHD4-GFP was efficiently coenriched in the presence of conjugated NEAT1_1, which also impaired the enrichment of endogenously biotinylated proteins. Interestingly, the protein domains that mediate binding of CHD4 to RNA have recently been mapped to the N-terminal part of CHD4 in *D. melanogaster* and human cells(74). Thus, we created the N-terminal deletion mutant ΔN-CHD4-GFP (Fig. S6A). We expressed both full length and ΔN-CHD4-GFP in HEK293 cells and compared the efficacy of binding to *in vitro* transcribed NEAT1_1 (Fig. 5E; Fig. S6B). Indeed, CHD4-GFP bound NEAT1_1 ∼2-fold stronger than the mutant. Of note, we also expressed CHD4-GFP variants in cells and noticed mislocalization of ΔN-CHD4-GFP to the cytoplasm irrespective of etoposide treatment (Fig. S6C). Thus, we did not attempt *in vivo* binding assays with CHD4-GFP variants. We conclude that CHD4 binds recombinant NEAT1_1 via its N-terminal part *in vitro* and associates with NEAT1 *in vivo*.

**Figure 5.**
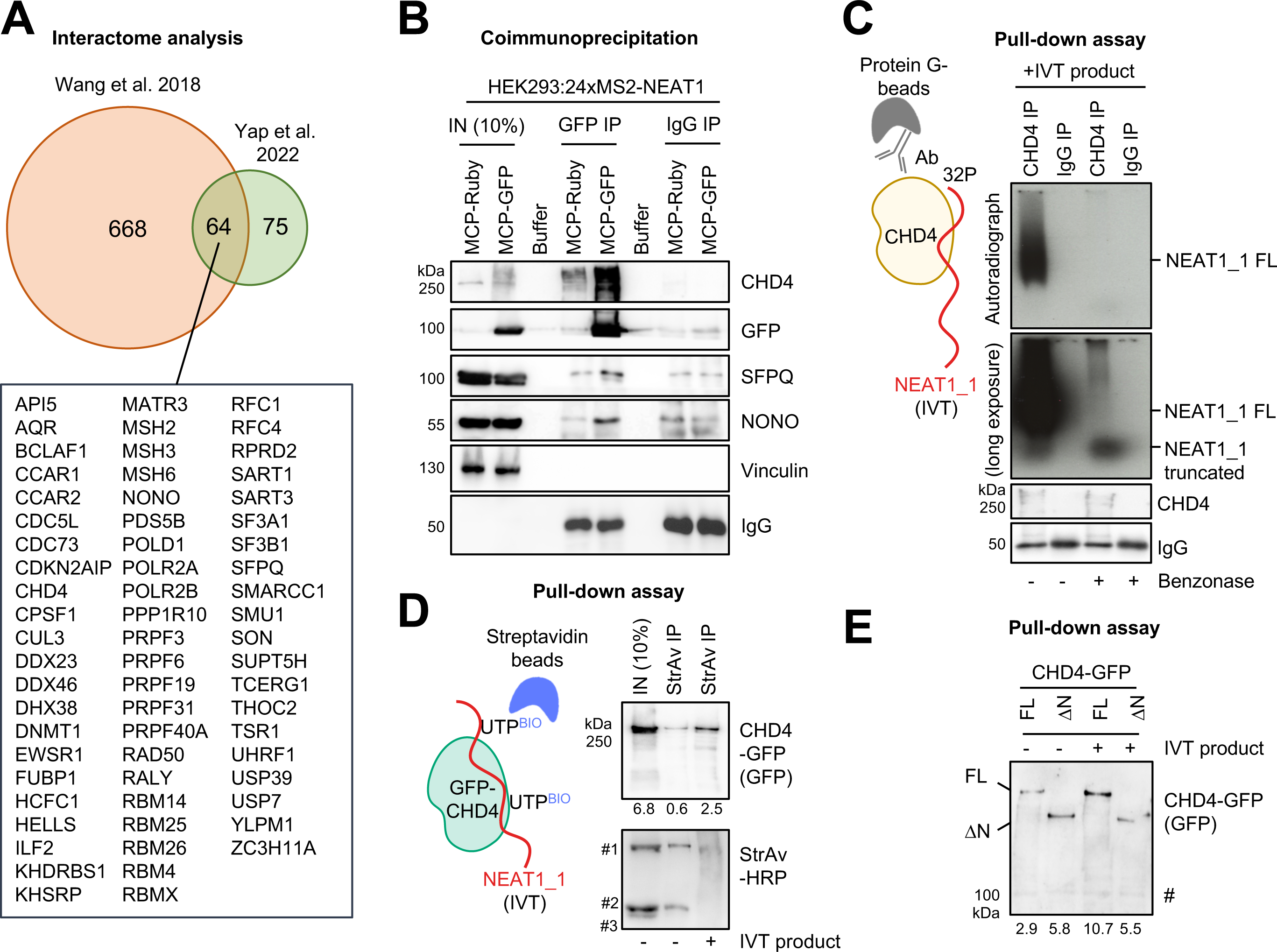
NEAT1 associates with CHD4 *in vivo* and *in vitro*. (**A**) Interactome analysis identifies 64 potential interactors of NEAT1. (**B**) Immunoblots detecting the chromodomain helicase DNA binding protein 4 (CHD4), GFP, SFPQ and NONO in whole cell lysates of MCP-GFP-expressing HEK293:24xMS2-NEAT1 cells (IN) or upon immunoselection with GFP antibody. Expression of MCP-Ruby and immunoselection with immunoglobulin G (IgG), controls. (**C**) Scheme of pull-down assay (left) and autoradiographic detection of 32P-γ-ATP end-labeled (32P) NEAT1_1 *in vitro* transcription (IVT) full length (FL) product upon immunoprecipitation (IP) with immobilized CHD4 from HEK293 whole cell lysates (right). Immunoblots, loading controls. (**D**) Scheme of pull-down assay (left) and detection of full length GFP-tagged CHD4 (CHD4-GFP) or endogenously biotinylated proteins (#1-3) by immunoblots upon ectopic expression in HEK293 cells and immunoprecipitation from whole cell lysates (IN) with biotin (BIO)-labelled and immobilized NEAT1_1 *in vitro* transcription (IVT) product. #1, Pyruvate carboxylase; #2, Propionyl-CoA carboxylase; #3, β-methylcrotonyl-CoA carboxylase. (**E**) Pull-down assay as in (D), but upon expression of full length CHD4-GFP (FL) or N-terminal mutant CHD4-GFP (ΔN). #, non-specific.

### The DNA damage response triggers the release of CHD4 from NEAT1 to promote chromatin occupancy

Chromatin-associated NEAT1 scaffolds the recruitment of some histone modifying enzymes and chromatin-remodelling factors(75, 76). Since NEAT1 interacts with CHD4, we asked if this association is altered by DNA damage. Using our HEK293:24xMS2-NEAT1 system, we repeated MS2-NEAT1 immunoselection from cells incubated in the presence or absence of etoposide or after preincubation with STM2457. Indeed, etoposide incubation largely impaired the association of CHD4 with MS2-NEAT1, but did not severely alter the expression levels of CHD4 (Fig. S7A). *Vice versa*, we incubated lysates of U2OS cells that were cultured in the absence or presence of etoposide with antibodies against CHD4 or the nucleolar RBP Fibrillarin and determined the amount of coimmunoprecipitating NEAT1 by RT-qPCR (Fig. 6A). Again, the association of NEAT1 to CHD4 was partially impaired by etoposide treatment, and not detectable by the Fibrillarin antibody. To test for DNA damage-responsive alterations in CHD4 RNA binding more globally, we performed sucrose gradients with lysates from cells that were incubated in the absence or presence of etoposide or pre-treated with STM2457. We observed a strong etoposide-responsive decrease of CHD4 in the higher molecular weight fractions 16 and 17, which was sensitive to STM2457 preincubation and accompanied by a relative decrease of NEAT1 in fractions 16 and 17 (Figs. 6B, 6C). The integrity of the assay was monitored by RNase digestion and probing for Vinculin and the *bona fide* RBP Nucleophosmin 1. The latter strongly shifted to lower molecular weight fractions in RNase-treated samples (Fig. S7B). Finally, we wished to test if CHD4 accumulates on broken chromatin in a METTL3-dependent manner. We performed CUT&RUN-seq. Assessing 80 accessible AsiSI cleavage sites, we found prominent occupancy of CHD4 around AsiSI-induced DSBs, which was partially impaired by STM2457 preincubation (Fig. 6D). When stratifying for promoter- and non-promoter-associated AsiSI sites, the CHD4 phenotype could also be observed at 63 non-promoter-associated AsiSI sites, which display poor NEAT1 occupancy in CHART-seq. In striking contrast, STM2457 preincubation partially reversed the phenotype at 17 promoter-associated AsiSI sites, which display high NEAT1 occupancy in CHART-seq. Visual inspection of browser tracks and manual ChIP for CHD4 at DS1 confirmed our analysis (Fig. 6E; Figs. S7C, S7D). Together, we conclude that METTL3 is required to release CHD4 from NEAT1 and promotes the spreading of CHD4 to DSB that are not occupied by NEAT1.

**Figure 6.**
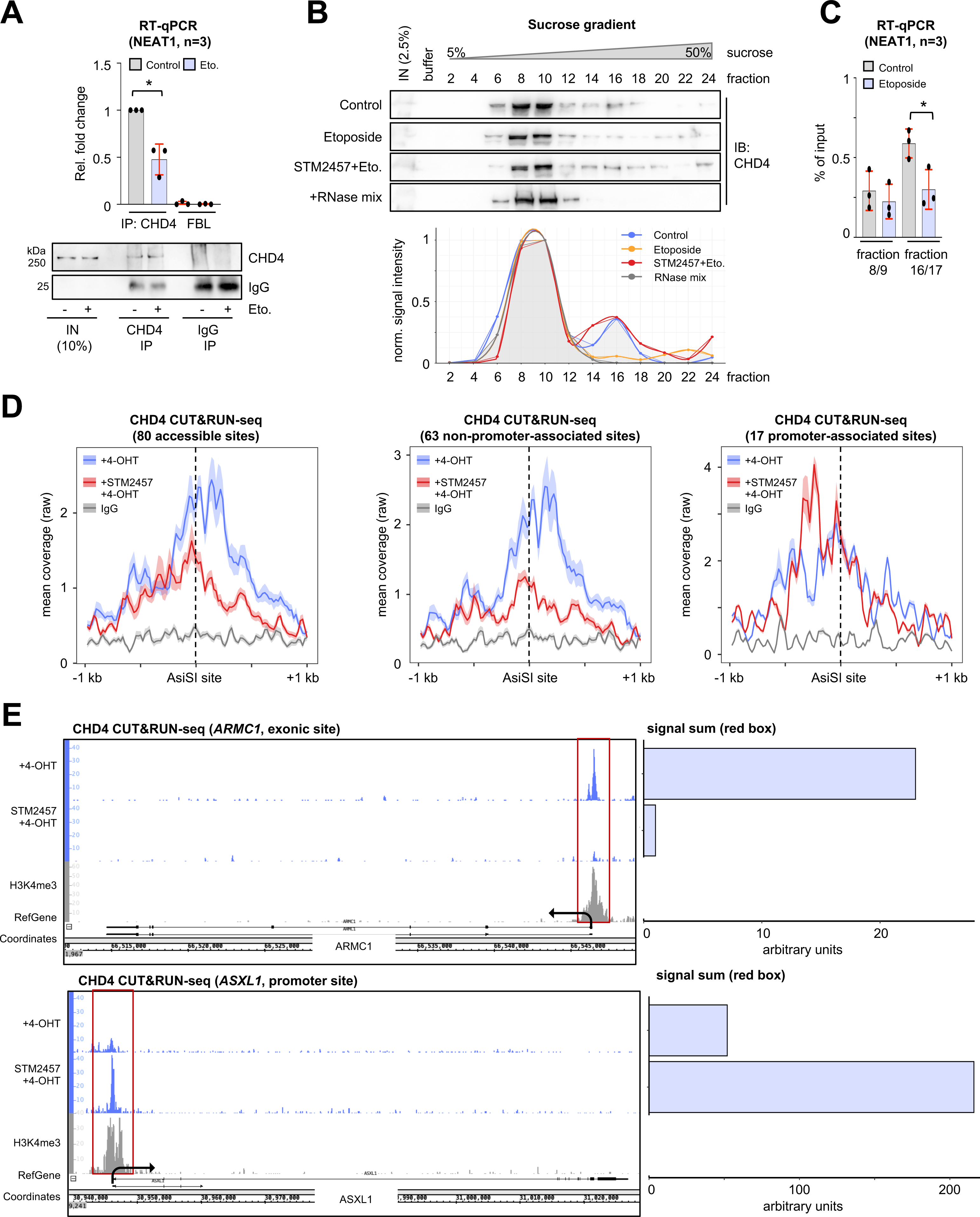
DNA damage reduces the association of CHD4 with NEAT1 in U2OS cells. (**A**) RT-qPCR assessing NEAT1 levels (top) and immunoblots detecting CHD4 in whole cell lysates (IN) and upon immunoprecipitation with selective antibodies. FBL (Fibrillarin) and IgG, controls. (**B**) Immunoblots (top) and quantitation (bottom) of CHD4 upon sucrose gradient fractionation. IN, input. (**C**) RT-qPCR assessing NEAT1 levels from sucrose gradient fractions. (**D**) CHD4 CUT&RUN-seq at AsiSI sites (dashed line). (**E**) Browser tracks (left) and quantitation (right) of CHD4 CUT&RUN-seq. Red box, region of interest; arrowhead, transcription start site; histone H3 lys-4 tri-methylation (H3K4me3), control. */**, p-value <0.05/ <0.001; two-tailed t-test. Error bar, mean ±SD. n=number of biological replicates.

## DISCUSSION

We describe NEAT1 as direct regulator of the DDR. NEAT1 undergoes structural rearrangement and accumulates at DSBs upon METTL3-dependent methylation to promote the release of CHD4 from NEAT1, which facilitates the efficient recognition and repair of a subset of DSBs (Fig. 7). It is important to clarify that we neither claim that METTL3 is solely targeting NEAT1 nor that NEAT1 is the sole regulator of CHD4 in DSBR. METTL3 has widespread roles in regulating RNA metabolism upon DNA damage, which includes the turnover of R-loops and the amplification of p53-signalling(55, 58). Likewise, the recruitment of CHD4 to broken chromatin is well-documented and, for instance, requires interaction with sequence-specific DNA binding proteins(77, 78). Intriguingly, CHD4 has also been described as binder of pre-mRNA transcripts(74). In this scenario, nascent pre-mRNA synthesis shields CHD4 from actively-transcribed promoters and thus maintains RNAPII activity. We observed that the inhibition of METTL3 diminished the occupancy of CHD4 at the bulk of AsiSI-induced DSBs. Thus, it is tempting to speculate that CHD4 binding to NEAT1 provides an additional layer to regulate promoter activity and that the release of CHD4 from NEAT1 upon DNA damage-responsive METTL3 activation may inhibit RNAPII activity at promoter-associated DSBs, which is a prerequisite for efficient DSBR(6, 7). It will be important to determine the impact of NEAT1 depletion on RNAPII activity in context of DSBR in the future.

**Figure 7.**
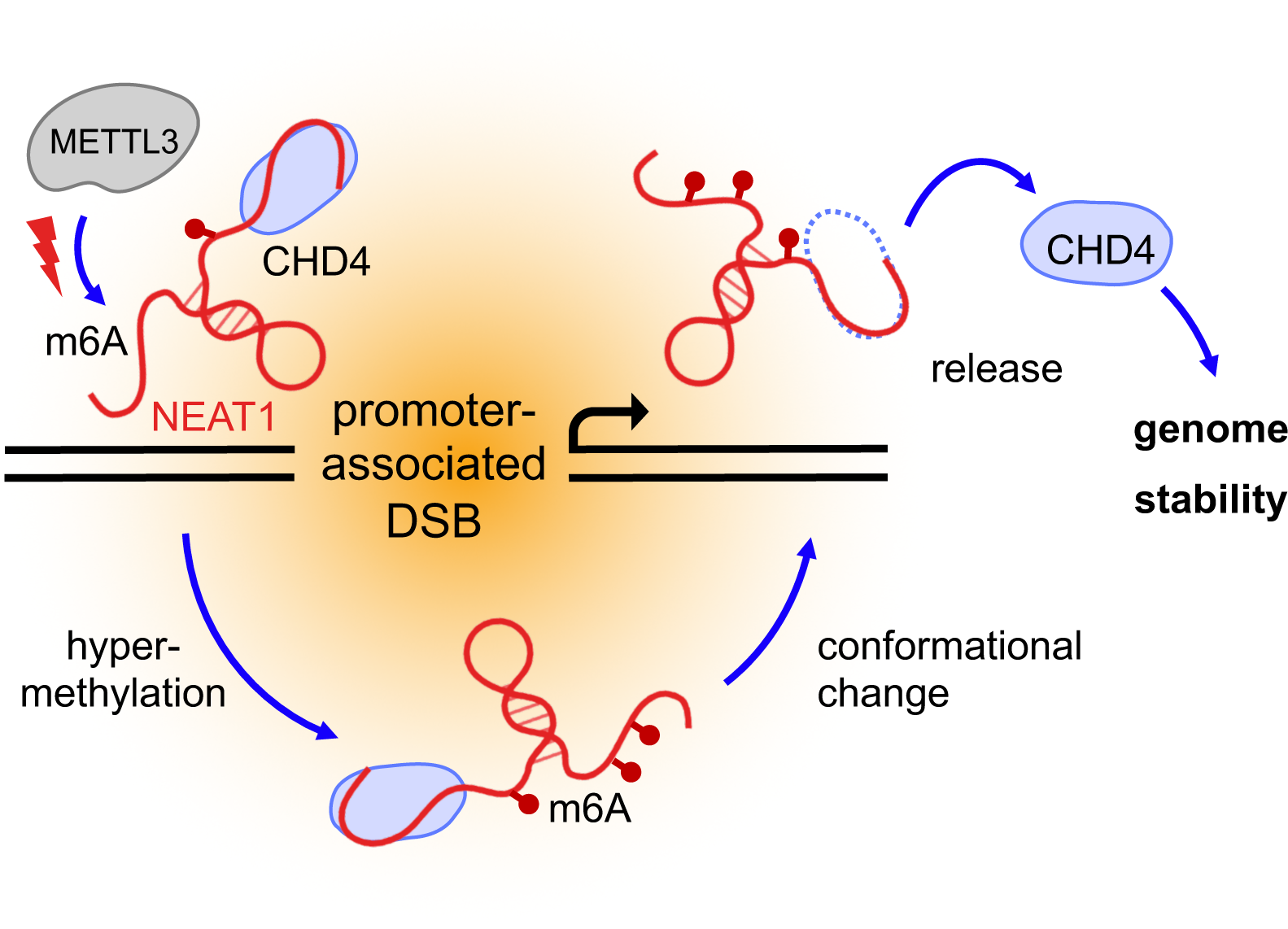
Model illustrating our findings. See main text for details.

A recent study observed prominent reduction of DDR factor expression upon NEAT1 depletion in leukaemia cells, which was accompanied with elevated levels of γH2A.X and endogenous DNA damage(45). Thus, our observation that elevated DNA damage in NEAT1-deficient cells is accompanied by diminished, rather than elevated induction of DSB signalling markers, such as the increase in γH2A.X levels and the formation of 53BP1-positive foci, is intriguing and may seem counterintuitive at first glance. Moreover, we observed prominent defects in genome stability only upon combining NEAT1 depletion with etoposide treatment. These apparent discrepancies may, at least in part, be explained by the different cellular systems used or due to differences in the expression level and depletion efficacies of NEAT1 itself. As our data have been obtained mostly under pulse-chase conditions, we postulate that NEAT1-deficient cells comprise defects in early and acute steps of DSB signalling. In line with this, we observed a modest accumulation of H2BK120ac marks, a DSB marker upstream of γH2A.X, NEAT1-deficient cells. This suggests a supportive rather than essential role for NEAT1 in genome stability. The diminished DSB signalling observed in NEAT1-deficient cells may be rather transient and may not fully translate into differences in the activation of downstream DSBR factors, which has also been concluded for other modes of RNA-dependent DSBR(13, 23).

Why do γH2A.X- and 53BP1-positive foci not form efficiently in NEAT1-deficient cells? Firstly, one or more components of the DSB signalling cascade may not be sufficiently expressed in NEAT1-deficient cells, which may prevent the efficient formation of γH2A.X- and 53BP1-positive foci. Studies in mice and murine cells suggest that the knockout NEAT1 provokes global changes in gene expression, which drives neoplastic cell growth in murine embryonic fibroblasts, but does not seem to be required for stress-induced cell cycle arrest or apoptosis(39). We did not observe severe alterations in the cell cycle distribution upon NEAT1 depletion and no obvious impact on the expression of METTL3 or CHD4. However, it remains to be determined to what extent the multi-layered post-transcriptional gene regulatory roles of NEAT1, which include pre-mRNA editing, miRNA biogenesis and translational control(34, 79–81), modulate our phenotypes. Secondly, and perhaps more appealing, NEAT1 may comprise an intrinsic property to promote DSB condensation. Arguing for the latter, we indeed observed the formation of γH2A.X- and 53BP1-positive foci in NEAT1-deficient cells, albeit to a lesser degree. Intriguingly, functional domains of NEAT1 have been identified in the middle domain of NEAT1_2, which promote the formation and condensation of nuclear paraspeckles via phase-separation(82). Since 53BP1 and other early recruiting DSB factors utilize DNA damage-induced transcripts for the efficient recruitment to DSBs(83–85), it appears possible that the intrinsic phase-separating property of NEAT1_2 is hijacked by the DDR to foster foci formation and thereby promote DSB condensation.

Our model suggests that NEAT1 engages at a subset of promoter-associated DSBs to modulate CHD4. In line with our model, recent ATAC-seq data from human colorectal cancer cells suggest that NEAT1 is indeed required for chromatin remodelling in response to genotoxic stress(38). However, whether NEAT1 engages in DSB signalling exclusively on chromatin or also in context of paraspeckles remains unclear. Whilst our CHART data suggest DNA damage-induced association of NEAT1 with a subset of DSBs, a rather modest colocalization of NEAT1 with 53BP1-positive DSBs foci could be observed in our RNA-FISH data, albeit displayed as non-staked confocal sections. Interestingly, the core paraspeckle components NONO, SFPQ and PSCP1 also occupy chromatin at a subset of protein-coding gene promoters in unperturbed cells(33). Moreover, the DDR triggers mobility of DSBs and formation of DNA damage-induced higher-order chromatin structures(86). Thus, it seems likely that only a fraction of the NEAT1 pool is amenable to the DDR and that a subset of DSBs may cluster in specialised subnuclear domains, including paraspeckles.

Of note, our data do not allocate the repair-promoting function of NEAT1 to a distinct isoform, as we used pooled probe set and oligonucleotides that cover both NEAT1_1 and NEAT1_2 for hybridization in most cases. This is partially due to technical constraints in sensitivity and also due to the biology of NEAT1, which is mostly accessible for hybridization studies in the regions that we were targeting, as determined by RNase H mapping, previously(60). Nevertheless, our *in vitro* data indicate that the short isoform is necessary and sufficient for at least some of the observed phenotypes.

Overall, our study describes a novel RNA-mediated DDR pathway and a new function for NEAT1 in the regulation of DSBR. Given that NEAT1 is highly abundant in many tumours, the validation of the genome-protective pathway in primary cancer material promises to be a powerful approach to understand the role of NEAT1 in patients and could pave the way for novel, RNA-centric therapeutic approaches. Current approaches to interfere with NEAT1 function in cancer cells, for instance, include the application of ASOs(87). This non-genotoxic strategy could be combined with chemotherapy to hypersensitize cancer cells to genotoxic drugs in the future.

### DATA AVAILABILITY

Sequencing data are available at the gene expression omnibus under the accession number GEO:GSEpending (reviewer token: pending), and are viewable on the integrated genome browser (IGB) or other suitable genome browsers. Further information and requests for resources and reagents should be directed to the corresponding author.

## ACKNOWLEDGMENTS

We acknowledge Elmar Wolf, Mathias Munschauer and Markus Diefenbacher for feedback and discussions. We thank Christian Janzen, Juliane Müller, Giacomo Cossa, Carsten Ade, Lea Boten, An Binh Nguyen and Tobias Roth for excellent technical support, and all members of the department for sharing reagents and collegial atmosphere. We apologize to authors whose work could not be cited due to limitations.

Author contributions: Conceptualization, V.M. and K.B.; Methodology, V.M., B.T., A.-S.G.-B., P.B., P.P., W.S., P.G., D.P. and K.B.; Investigation, V.M., B.T., A.-S.G.-B., W.S. and K.B.; Formal Analysis, V.M., B.T., W.S., P.B., P.G., D.P. and K.B.; Writing – Original Draft, V.M. and K.B.; Writing – Review & Editing, V.M. and K.B.; Funding Acquisition, M.E., T.G., R.P.S. and K.B.; Supervision, M.E., T.G., R.P.S. and K.B.

## SUPPLEMENTARY DATA

Supplementary Data are available online.

