## Supplementary Information for "NEAT1 promotes genome stability via m6A methylation-dependent regulation of CHD4"

**Supplementary information contains 7 supplementary figures and 6 supplementary tables.**

### Figure S1

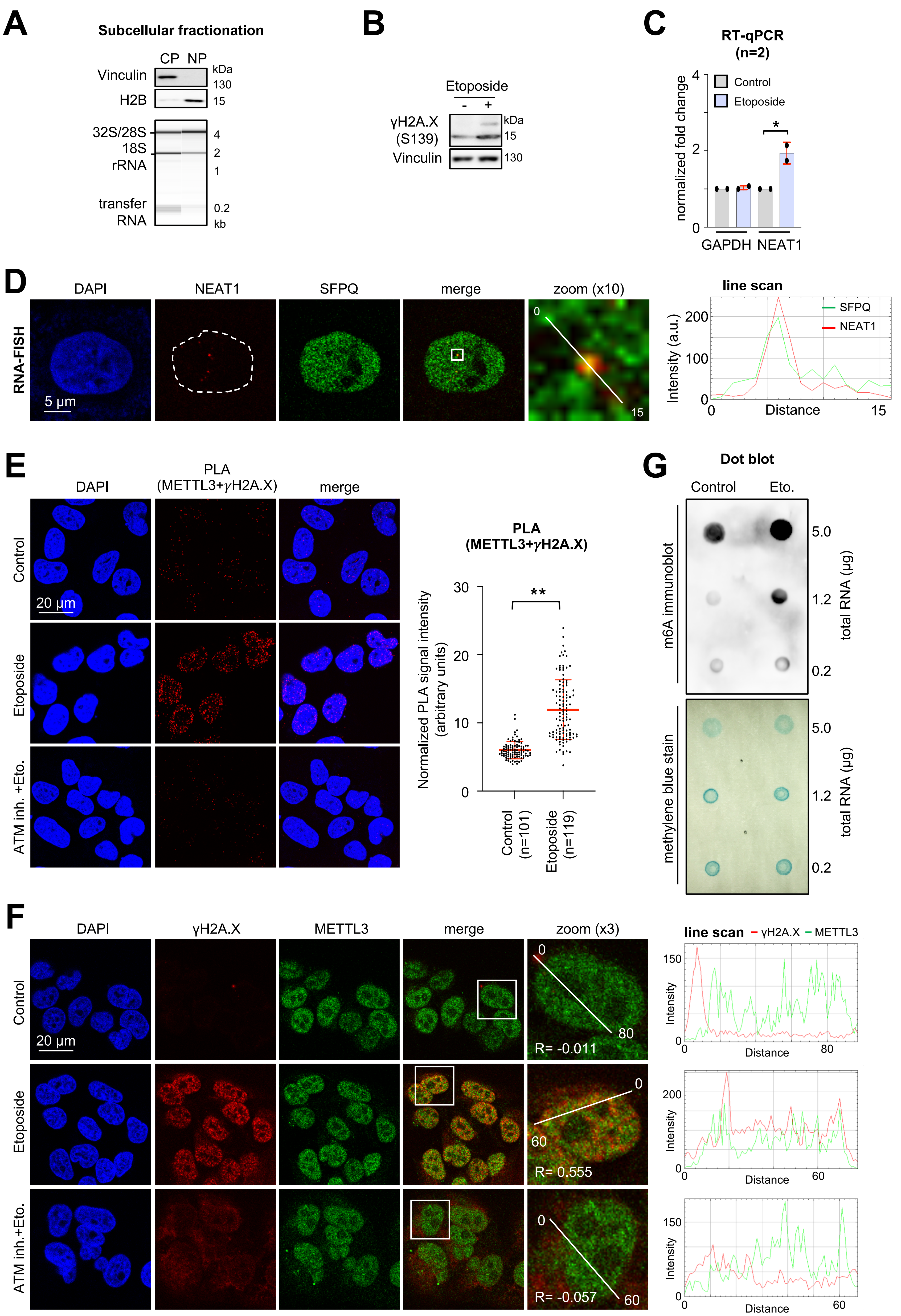

**Figure S1.** Quality controls for subcellular fractionation, induction of DNA damage, and quantitation and imaging of NEAT1. (**A**) Immunoblots detecting Vinculin and histone H2B (top) and Fragment Analyzer gel detecting ribosomal (r)RNA and transfer RNA (bottom) upon subcellular fractionation into cytoplasm (CP) and nucleoplasm (NP) of U2OS cells. (**B**) Immunoblots detecting serine-139 phosphorylated H2A.X (γH2A.X) in U2OS cells. Vinculin, loading control. (**C**) RT-qPCR assessing NEAT1 levels in HEK293 cells. Glyceraldehyde-3-phosphate dehydrogenase (GAPDH), control. (**D**) Imaging (left) and line scan quantitation of colocalisation (right) of Quasar570-labelled RNA-FISH probe and SFPQ antibody signals. More than 80 cells were assessed. Broke white circle, nucleus; white box, zoom. (**E, F**) Imaging (left) and quantitation (right) of PLA signals (E) or indirect immunofluorescence signals (F) for METTL3/γH2A.X. Each dot represents one acquisition; white box, zoom; R, Pearson correlation coefficient. (**G**) Dot blot analysis of total RNA. Methylene blue stain, loading control. */**, p-value <0.05/ <0.001; two-tailed t-test. Error bar, mean ±SD. n=number of biological replicates or imaged cells. Representative images are shown.

### Figure S2

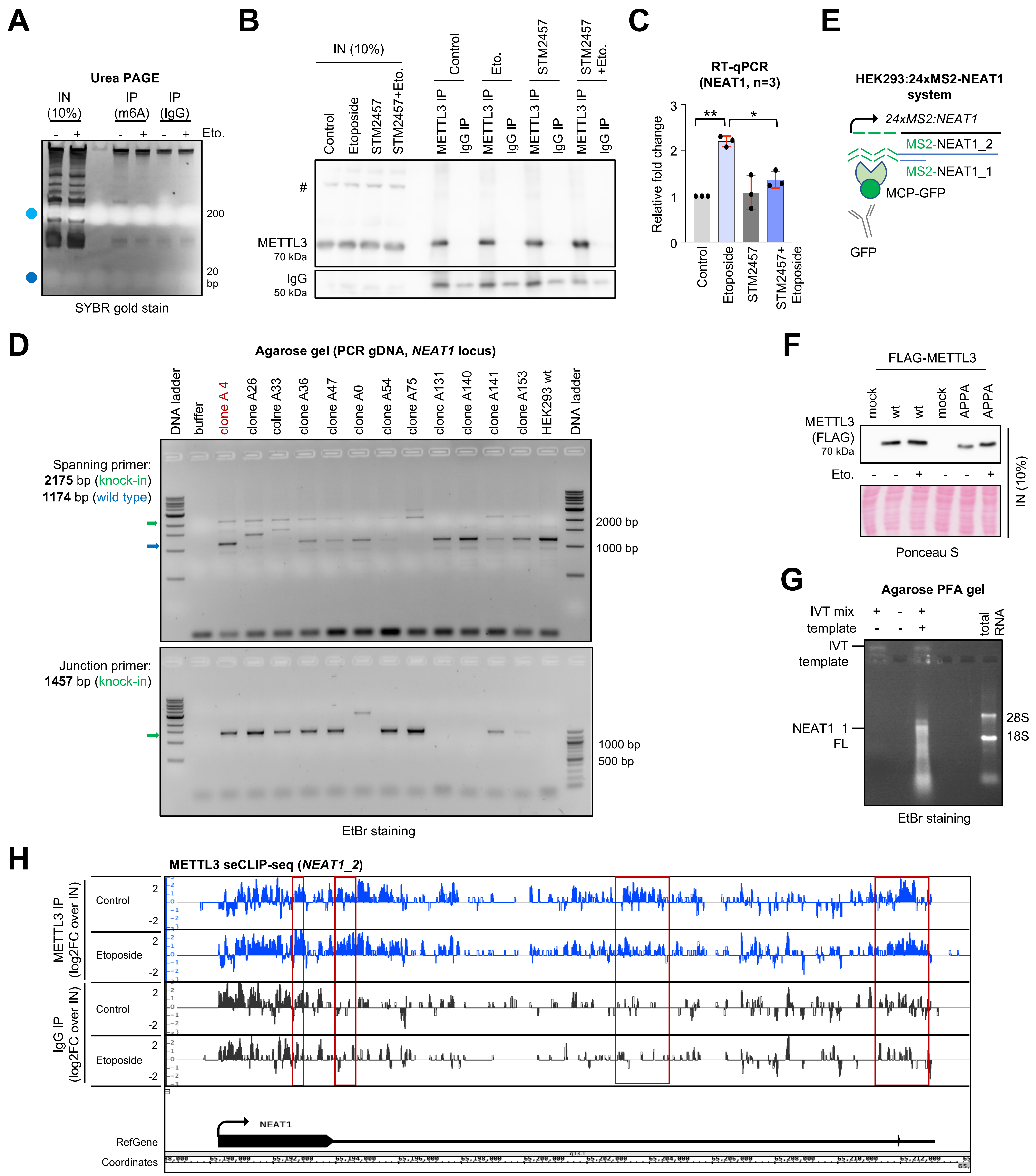

**Figure S2.** Quality controls for RIP assay, STM2457 treatment, CRISPR-tagging and *in vitro* transcription. (**A**) Urea PAGE gel displaying SYBR gold-stained RNA from U2OS whole cell lysate (IN) or upon immunoprecipitation (IP) with antibodies. Light/dark blue, xylene cyanol/bromophenol blue running front. (**B**) Immunoblot detecting METTL3 from U2OS whole cell lysates (IN) or upon immunoprecipitation (IP). IgG, control; #, non-specific. (**C**) RT-qPCR assessing NEAT1 levels in U2OS cells. (**D**) Agarose gel displaying ethidium bromide (EtBr)-stained PCR products from genomic (g)DNA of HEK293 CRISPR/Cas9 clones. Green/blue arrowhead, knock-in/wild type allele. (**E**) Scheme of the HEK293:24xMS2-NEAT1 system for immunoselection of MS2-tagged NEAT1. (**F**) Immunoblot displaying ectopically expressed FLAG-tagged METTL3 variants from whole cell lysates (IN). Ponceau S, loading control. (**G**) Agarose paraformaldehyde (PFA) gel displaying EtBr-stained *in vitro* transcription (IVT) product or total RNA from U2OS cells. 28S/18S, size marker. (**H**) Browser tracks of METTL3 seCLIP-seq reads at the *NEAT1_2* locus in U2OS cells. Read log2 fold-changes between merged seCLIP-seq duplicates and size-matched input are shown. Red box, region of binding; arrowhead, transcription start site. */**, p-value <0.05/ <0.001; two-tailed t-test. Error bar, mean ±SD. n=number of biological replicates.

### Figure S3

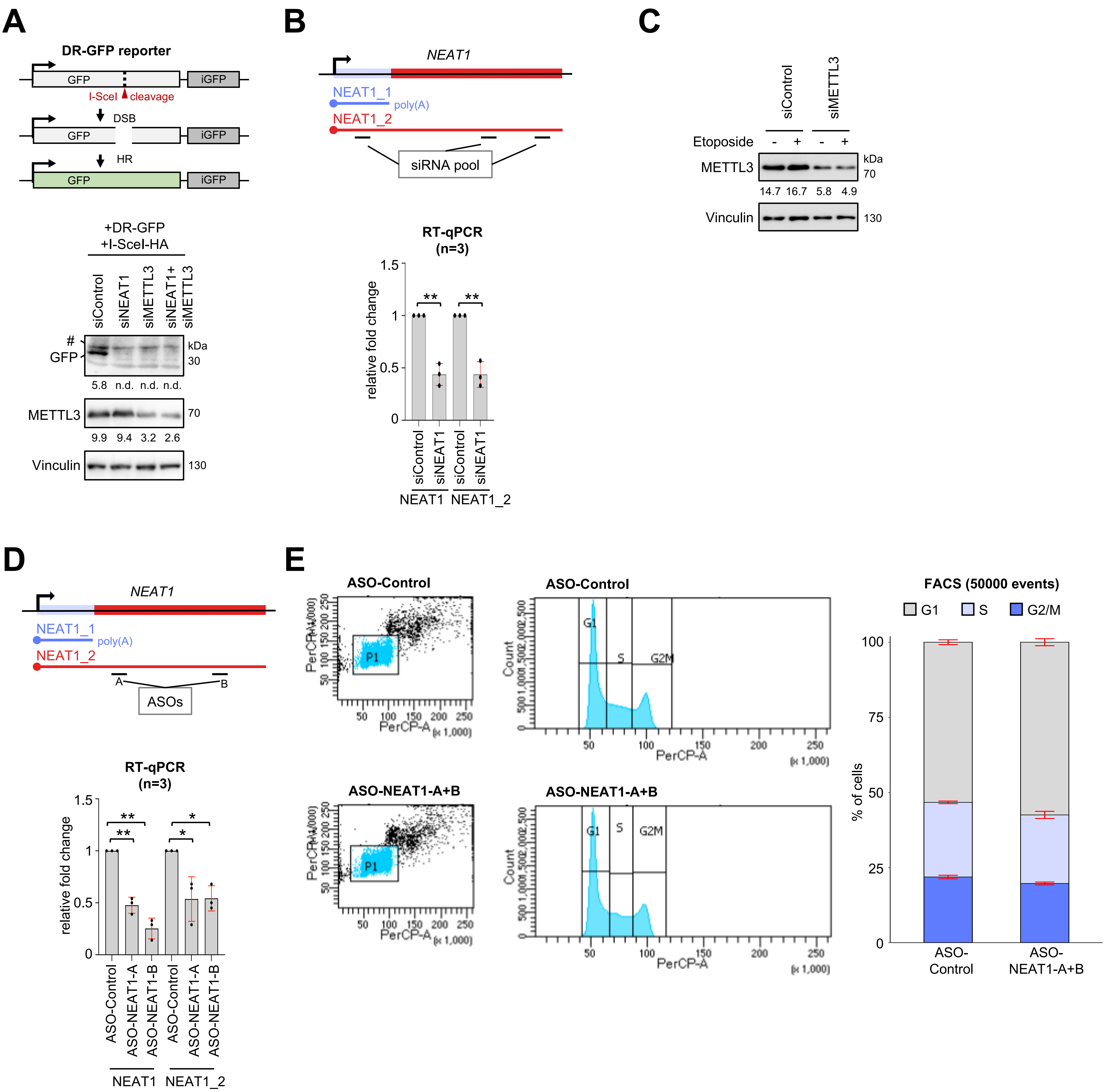

**Figure S3.** DSB repair reporter assay and quality controls for NEAT1 depletion. (**A**) Scheme of DR-GFP homologous recombination (HR) reporter containing I-SceI cleavage site (red arrowhead) and HR repair template (iGFP) (top) and immunoblots (bottom) detecting GFP and METTL3 levels upon plasmid and siRNA co-transfections in HEK293 cells. Vinculin, loading control; #, non-specific. (**B**) Scheme depicting landing sites for siRNA (top) and RT-qPCR assessing total NEAT1 and NEAT1_2 levels in U2OS cells (bottom). (**C**) Immunoblots detecting METTL3 upon siRNA transfection in U2OS cells. Vinculin, loading control. (**D**) Scheme depicting landing sites for antisense oligonucleotides (ASOs) A and B (top) and RT-qPCR assessing total NEAT1 and NEAT1_2 levels in U2OS cells (bottom).(**E**) Cell cycle analysis by FACS upon ASO transfection. Gating (P1) of propidium iodide-positive, viable, non-duplet cells (left) and stratification for cell cycle phase (right). PerCP-A/PerCR, forward/sideward scatter. A representative experiment is shown. */**, p-value <0.05/ <0.001; two-tailed t-test. Error bar, mean ±SD. n=number of biological replicates.

### Figure S4

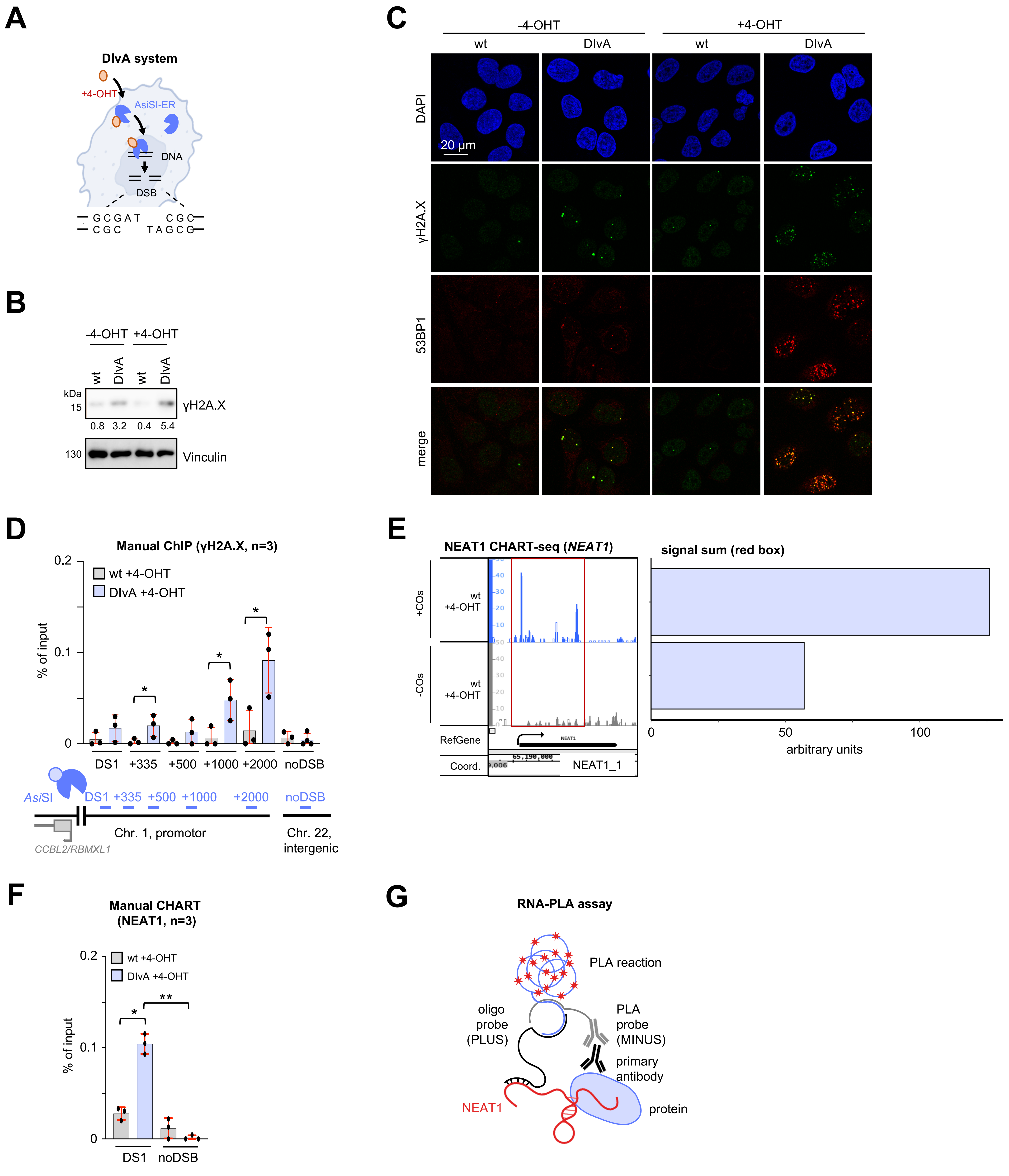

**Figure S4.** Quality controls for DIvA system and CHART assay. (**A**) Scheme of the DIvA U2OS system. 4-OHT, 4-hydroxytamoxifen. (**B**) Immunoblots detecting γH2A.X levels in wild type U2OS and DIvA cells. Vinculin, loading control. (**C**) Imaging detecting γH2A.X- and 53BP1-positive foci in wild type U2OS and DIvA cells. (**D**) Manual ChIP for γH2A.X at the AsiSI site DS1 in wild type U2OS and DIvA cells (top) and scheme of the AsiSI site DS1 (upstream of *RBMXL1/CCBL2* promoter). noDSB, non-restricted control. (**E**) NEAT1 CHART-seq browser tracks of *NEAT1* gene. COs, capturing oligos; red box, region of interest; arrowhead, transcription start site. (**F**) Manual CHART for NEAT1 at DS1 in wild type U2OS and DIvA cells. noDSB, non-restricted control. (**G**) Scheme for RNA-PLA assay. A selective antibody is combined with a DNA oligonucleotide that contains (i) a sequence complementarity with the RNA of interest, (ii) a linker sequence and (iii) a sequence that is utilized as PLUS probe during the PLA reaction. */**, p-value <0.05/ <0.001; two-tailed t-test. Error bar, mean ±SD. n=number of biological replicates. Representative images are shown.

### Figure S5

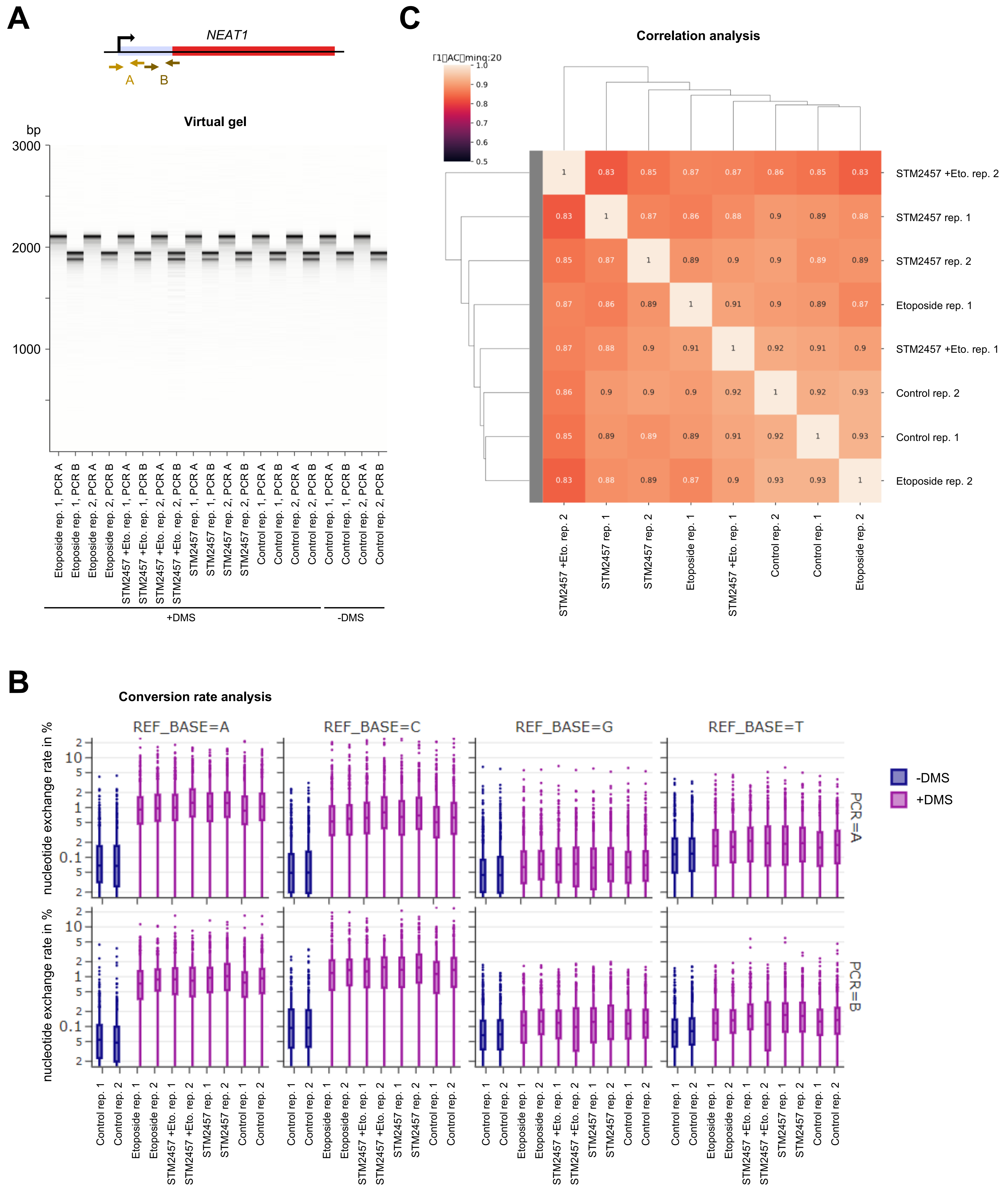

**Figure S5.** Quality controls for Nano-DMS-MaP-seq in HEK293 cells. (**A**) Primer locations for cDNA amplification of DMS-labelled NEAT1_1 (top) and virtual gel displaying 2 biological replicates of NEAT1_1 cDNA parts A and B. DMS, dimethyl sulfate. (**B**) Graphs displaying nucleotide exchange rates for 2 replicates of NEAT1_1 cDNA parts A and B upon Nanopore sequencing. (**C**) Correlation analysis of 2 biological replicates for each condition.

### Figure S6

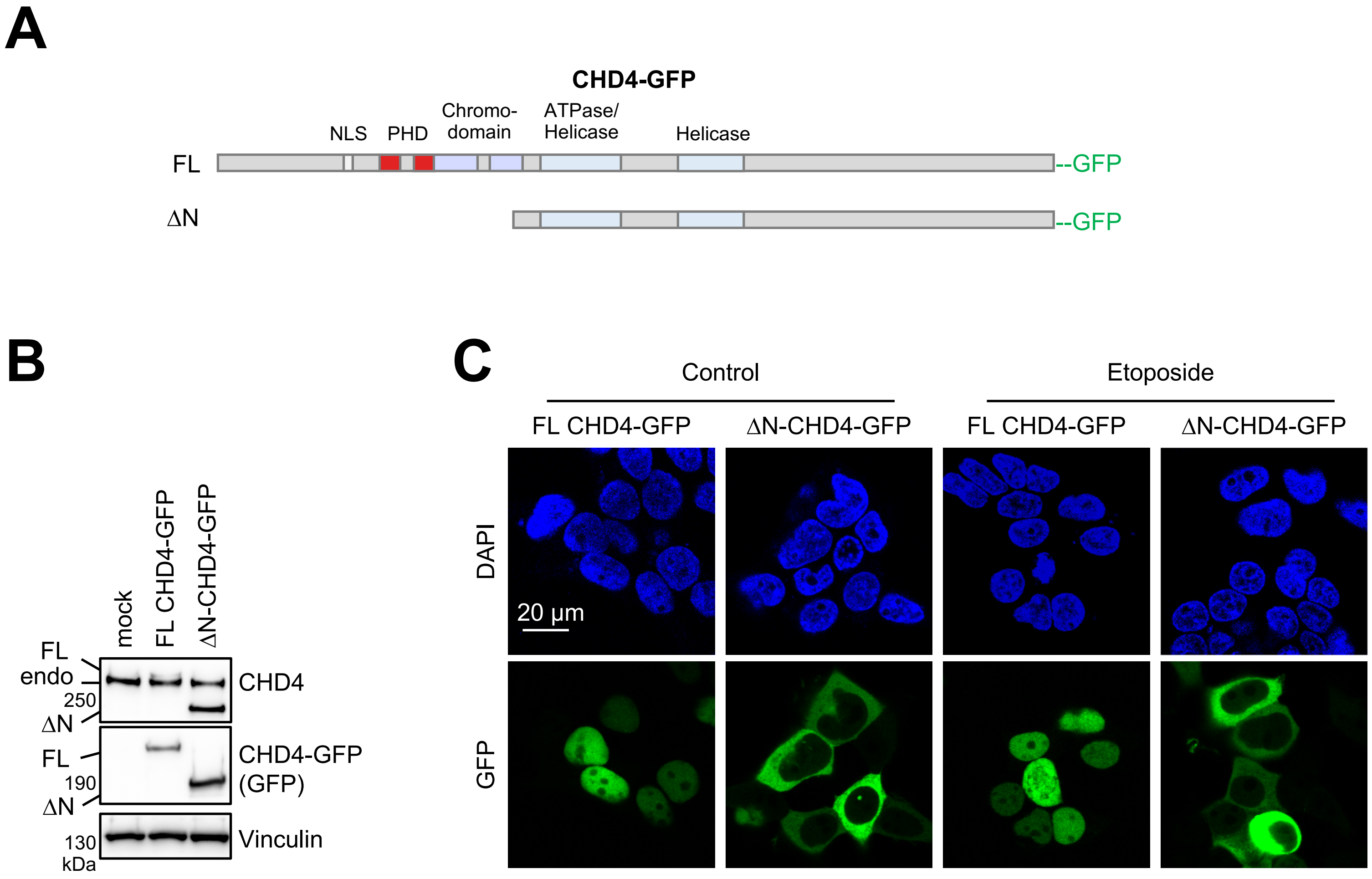

**Figure S6.** Quality controls for CHD4-GFP variants. (**A**) Scheme of CHD4-GFP variants. NLS, nuclear localisation signal; PHD, plant homeodomain; FL, full length; ΔN, N-terminal deletion mutant. (**B**) Immunoblots detecting endogenous CHD4 and recombinant CHD4-GFP variants. Vinculin, loading control. (**C**) Imaging of GFP signal upon ectopic expression of CHD4-GFP constructs in HEK293 cells. Representative images are shown.

### Figure S7

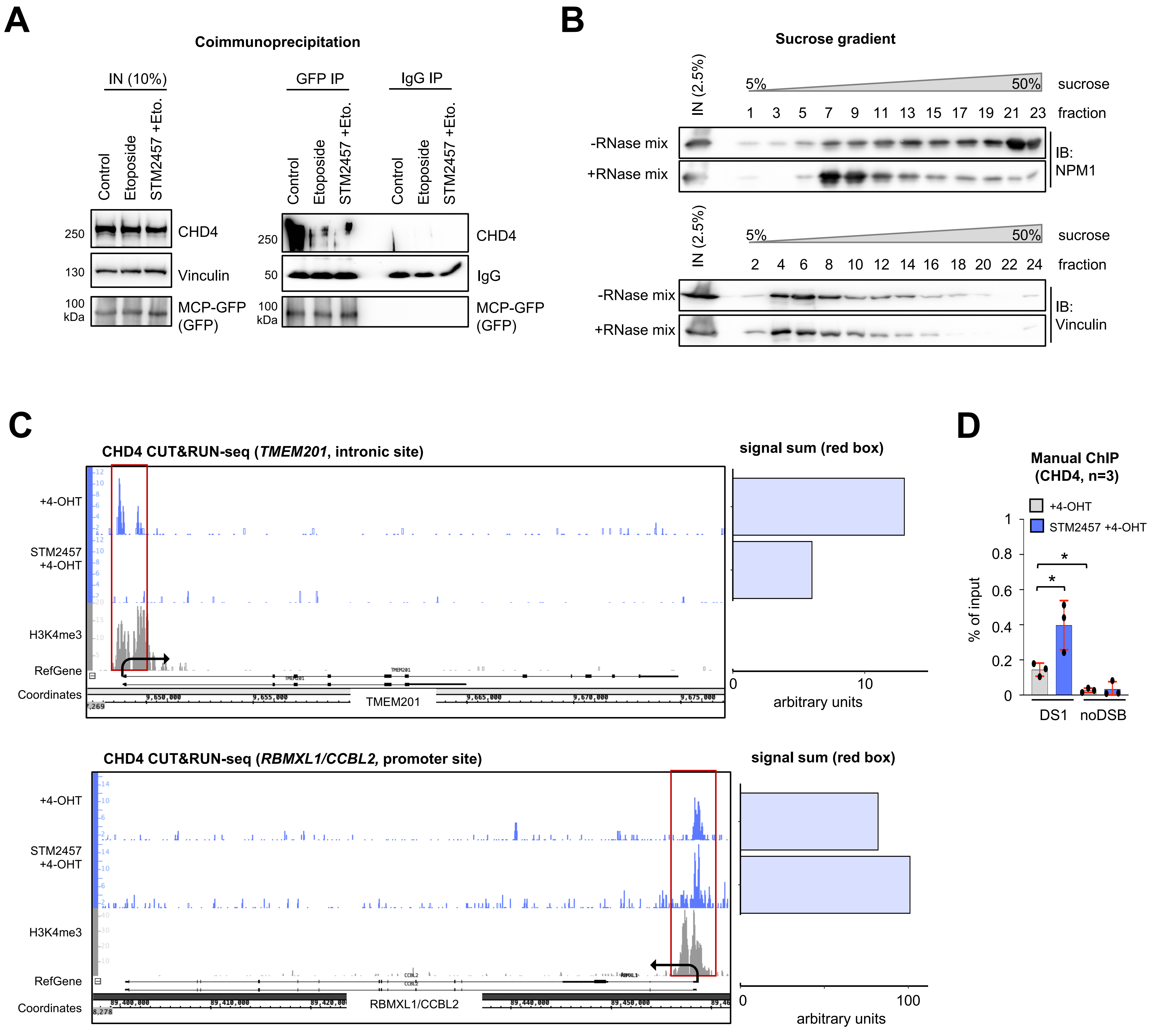

**Figure S7.** Quality controls for sucrose gradients and CUT&RUN-seq. (**A**) Immunoblots detecting CHD4 in whole cell lysates (IN) of HEK293:24xMS2-NEAT1cells that ectopically express MCP-GFP (left) and upon immunoselection with GFP antibodies (right). Vinculin and IgG, loading controls. (**B**) Immunoblots (IBs) detecting Nucleophosmin 1 (NPM1) (top) and Vinculin (bottom) signals upon sucrose gradient fractionation in U2OS cells. IN, input. (**C**) Browser tracks (left) and quantitation (right) of CHD4 CUT&RUN-seq in U2OS cells. Red box, region of interest; arrowhead, transcription start site; histone H3 lys-4 tri-methylation (H3K4me3), control. (**D**) Manual ChIP for CHD4 at AsiSI site DS1 in U2OS cells. noDSB, non-restricted control. */**, p-value <0.05/ <0.001; two-tailed t-test. Error bar, mean ±SD. n=number of biological replicates.

**Table S1.** siRNA used for RNA interference.

| **siRNA** | **Sequence (5’-3’)** | **Supplier, code** |
|---|---|---|
| siControl | pre-designed smart pool | Dharmacon, D-001810-01 |
| siMETTL3 | pre-designed smart pool<br><br>(siRNA-IDs: 132906, 132907, 132908) | Ambion, AM167008 |
| siNEAT1 | pre-designed smart pool<br><br>(siRNA-IDs: n272460, n341850, n509900) | Ambion, 4390815, 4392421 |

**Table S2.** ASOs used for transfections. All bases are DNA except ones preceded by 'm' like mU, which are 2'-hydroxy methylated RNA bases. *, phosphorothioate linkage. ASOs were custom made (IDT).

| **ASO** | **Sequence (5'-3')** |
|---|---|
| ASO-Control | mG*mC*mG*mU*mA*T*T*A*T*A*G*C*C*G*A*mU*mU*mA*mA*mC |
| ASO-NEAT1-A | mU*mG*mC*mG*mG*C*C*T*A*T*T*C*C*T*C*mC*mU*mG*mA*mC |
| ASO-NEAT1-B | mG*mC*mU*mG*mG*C*A*T*T*C*A*T*G*G*G*mC*mU*mC*mU*mG |

**Table S3.** Antibodies used in this study.

| **Primary antibody** | **Species** | **Supplier, code** |
|---|---|---|
| IgG control | rabbit | Proteintech, 30000-0-AP |
| Anti-NONO | rabbit | Proteintech, 11058-1-AP |
| Anti-SFPQ [EPR11874] | rabbit | Abcam, ab177149 |
| Anti-NPM1 [FC82291] | mouse | Abcam, ab10530 |
| Anti-phospho-histone H2A.X (S139) | rabbit | Cell Signaling, 2577 |
| Anti-phospho-histone H2A.X (S139) [JBW301] | mouse | Millipore, 05-636 |
| Anti-fibrillarin | rabbit | Abcam, ab5821 |
| Anti-vinculin | mouse | Sigma, V9131 |
| Anti-53BP1 | rabbit | Novus, NB100-304 |
| Anti-phospho-ATM/ATR substrate (S*Q) [D23H2/D69H5] | rabbit | Cell Signaling, 9607 |
| Anti-histone H2BK120ac | rabbit | Active Motif, 39120 |
| Anti-FLAG tag [M2] | mouse | Sigma, F-1804 |
| Anti-GFP tag | rabbit | Abcam, ab290 |
| Anti-histone H3K4me3 | rabbit | Abcam, ab8580 |
| Anti-METTL3 | rabbit | Proteintech, 15073-1-AP |
| Anti-m6A | mouse | Diagenode, C15200082-50 |
| Anti-NBS1 | rabbit | Novus, NB100-143SS |
| Anti-histone H2B | rabbit | Abcam, ab1790 |
| Anti-CHD4 | rabbit | Abcam, ab240640 |
| **Secondary antibody** | **Species** | **Supplier, code** |

| | | |
|---|---|---|
| HRP-linked IgG | mouse | Cytivia, NA931 |
| HRP-linked IgG | rabbit | Cytivia, GEHENA934 |
| Alexa Fluor 546-linked IgG | rabbit | Thermo, A10040 |
| Alexa Fluor 488-linked IgG | mouse | Thermo, A32766TR |
| Alexa Fluor 546-linked IgG | mouse | Thermo, A10036 |
| Alexa Fluor 488-linked IgG | rabbit | Thermo, A21206 |

**Table S4.** DNA probes used for RNA-PLA. Probes were custom made (Sigma).

| **Probe** | **Sequence (5'-3')** |
|---|---|
| NEAT1-sense-1 | GCACCCCCAGCCCGCCCCCTCGTCTCATCTAACTCAGTCATCTCTC AAAAAAAAAAAAAAAAAAAAATATGACAGAACTAGACACTCTT |
| NEAT1-sense-2 | TCTGTCCCCTGAAGCCCTGAGCTAGCCACTTCCTCCCCCACAACTA AAAAAAAAAAAAAAAAAAAAATATGACAGAACTAGACACTCTT |
| NEAT1-antisense-1 | GAGAGATGACTGAGTTAGATGAGACGAGGGGGCGGGCTGGGGGTGC AAAAAAAAAAAAAAAAAAAAATATGACAGAACTAGACACTCTT |
| NEAT1-antisense-2 | TAGTTGTGGGGGAGGAAGTGGCTAGCTCAGGGCTTCAGGGGACAGA AAAAAAAAAAAAAAAAAAAAATATGACAGAACTAGACACTCTT |

**Table S5.** Primer pairs used for RT-qPCR, manual ChIP and manual CHART. Primers were custom made (Sigma).

| **Primer** | **Sequence (5'-3')** |
|---|---|
| NEAT1-fwd | TTGACCAACGCTTTATTTTC |
| NEAT1-rev | TTACCAACAATACCGACTCC |
| NEAT1_2-fwd | GACTTCATTTCGAGTGATGG |
| NEAT1_2-rev | TTTCATCTGAACAGGGAATC |
| noDSB-fwd | ATTGGGTATCTGCGTCTAGTGAGG |
| noDSB-rev | GACTCAATTACATCCCTGCAGCT |
| DS1-fwd | GATTGGCTATGGGTGTGGAC |
| DS1-rev | CATCCTTGCAAACCAGTCCT |
| DS1+335-fwd | GAATCGGATGTATGCGACTGATC |
| DS1+335-rev | TTCCAAAGTTATTCCAACCCGAT |
| DS1+500-fwd | CCTGGATATGAGTTTGATCAGC |
| DS1+500-rev | CTCTCCTTTCGCTGACACTG |
| DS1+1000-fwd | AGGAATTGACTGCGGTGTTC |
| DS1+1000-rev | GGGGAGGAGGAAAGGTGTAG |
| DS1+2000-fwd | GCCATAACAGAGGGTGGAAA |
| DS1+2000-rev | AACTTTAGGATGGGGCTGCT |
| HPRT1-fwd | AGATGTGATGAAGGAGATGG |
| HPRT1-rev | AATAGCTCTTCAGTCTGATAAAATC |
| GAPDH-fwd | AACCTGCCAAATATGATGAC |
| GAPDH-rev | AGGAAATGAGCTTGACAAAG |

**Table S6.** DNA probes used as capturing oligos (COs) for CHART. Probes were custom made (IDT). iSp18, internal spacer; BioTEG, N,N'-Bisbiotin-tetra(ethylene glycol)-diamine.

| **CO** | **Sequence (5'-3')** |
|---|---|
| NEAT1-1 | CTAGCCACTTCCTCCCCCACAA/iSp18/BioTEG |
| NEAT1-2 | TGTCTGTCCCCTGAAGCCCTG/iSp18/BioTEG |
| NEAT1-3 | TTCCTTCTCGCACCCCCAGC/iSp18/BioTEG |
| NEAT1-4 | CTTACAAGGCCTCAGAAATG/iSp18/BioTEG |
| NEAT1-5 | GCTAGTGCTAAGGAGCTCAG/iSp18/BioTEG |
| NEAT1-6 | ATGAAGTCAGACCAGCAAAC/iSp18/BioTEG |
